# Modernized Tools for Streamlined Genetic Manipulation of Wild and Diverse Symbiotic Bacteria

**DOI:** 10.1101/202861

**Authors:** Travis J. Wiles, Elena S. Wall, Brandon H. Schlomann, Edouard A. Hay, Raghuveer Parthasarathy, Karen Guillemin

## Abstract

The capacity to associate symbiotic bacteria with vital aspects of plant and animal biology is outpacing our understanding of the mechanisms shaping these interactions. A major barrier to mechanistic studies is the paucity of tools for genetically manipulating wild and diverse bacterial isolates. Solving this problem is crucial to elucidating the cellular and molecular rules that govern symbiotic relationships and ultimately harnessing them for agricultural and biomedical applications. Therefore, we constructed a series of vectors that expedite genetic knock-in and knock-out procedures across a range of bacterial lineages. This was accomplished by developing strategies for domestication-free bacterial conjugation, designing plasmids with customizable features, and streamlining allelic exchange using visual markers of homologous recombination. These tools enabled a comparative study based on live imaging of diverse bacterial symbionts native to the zebrafish intestine, with which we discovered heterogeneous colonization patterns and a striking correlation between bacterial population biogeography and cellular behavior.

## INTRODUCTION

High-throughput metagenomic sequencing has exposed the previously unseen diversity of symbiotic bacteria that live in close contact with plants and animals throughout the biosphere^1,2^. Associations are being made at breakneck speed between the membership and activity of resident bacteria and the health, development, and evolution of their hosts^3–7^. However, the cataloging of symbiotic relationships—whether mutualistic, commensal, or pathogenic—is vastly outpacing their cellular and molecular interrogation^8,9^. Elucidating the mechanisms by which symbiotic bacteria live and interact with each other and their hosts will inform how they can be harnessed for agricultural and biomedical applications^2,10–12^.

Characterizing the biology of symbiotic bacteria requires methods for precisely manipulating their genomes. For example, stable chromosomal insertion of genes encoding fluorescent proteins allows cellular behaviors and interactions to be directly observed within native host-associated environments^13,14^. Additionally, gene deletion and complementation studies are essential for rigorously dissecting the genetic pathways that control specific phenotypes^15^. Such knock-in and knock-out technologies have long been mainstays in microbiology labs working with entrenched model organisms like *E. coli*, but established genetic approaches are often inadequate for manipulating wild and novel species or strains^16^. This is largely because legacy protocols can involve cumbersome and outdated procedures that are difficult to use across lineages. Consequently, the in-depth study of most symbiotic bacteria remains out of reach.

A major bottleneck within the field of symbiosis research is that locating appropriate genetic tools and methods or developing them de novo are arduous and time-consuming tasks. This problem is especially burdensome for investigators aiming to manipulate multiple bacterial lineages derived from complex communities. To overcome these barriers, we have employed the zebrafish intestinal microbiota as a source of wild and diverse symbiotic bacteria^17^—which includes representatives of the *Vibrio*, *Aeromonas*, *Pseudomonas*, *Acinetobacter*, *Enterobacter*, and *Plesiomonas* genera—to develop and test streamlined tools and methods for bacterial genetic manipulation. We identified three main deficiencies inherent to current genetic approaches that if resolved, will immediately improve the genetic tractability of many bacteria. First, although conjugation is a robust and reliable method for delivering DNA into bacteria, strategies for selecting individual cells carrying the transferred DNA are not broadly compatible between different lineages and sometimes rely on deleterious domestication steps. Second, most vectors used for making genetic manipulations are not readily customized, which restricts their versatility and prevents further innovation. And third, techniques for generating chromosomal modifications via allelic exchange often depend on specific selection conditions that can vary between bacterial lineages and are difficult to troubleshoot when they fail. To address these shortcomings, we rationally designed a centralized set of genetic engineering vectors with new and updated functionalities. For DNA delivery, we developed alternative schemes for post-conjugation counterselection that avoid initial domestication of engineered bacteria, thereby preserving their natural physiology and behavior. For customization, we designed gene expression scaffolds with interchangeable sequence elements that can be tailored to different bacterial genomes, and with these produced a variety of ready-made vectors for fluorescently tagging bacteria. Moreover, an extensive collection of marked zebrafish intestinal symbionts was generated during this work that will accelerate research in the growing zebrafish-microbiota community. Lastly, we devised a means of visually following homologous recombination events during allelic exchange protocols for more tractable generation of markerless chromosomal alterations.

To demonstrate the potential of these modernized tools to uncover new aspects of host-microbe interactions, we examined the colonization patterns of several bacterial symbionts native to the larval zebrafish intestine by light sheet fluorescence microscopy^18^. The intestinal microbiota is an especially important target for exploration because of its impact on host health and disease; however, its phylogenetic diversity and concealed location make it difficult to investigate in situ by conventional techniques. Unexpectedly, live imaging of bacterial symbiont behavior within larval zebrafish revealed that genome sequences and in vitro-based phenotypes were poor predictors of whether a given bacterium exhibits free- swimming motility in vivo. Most strikingly, we also discovered a general relationship between the growth mode of individual bacteria and their overall biogeography; namely, the average location of a population along the intestinal tract is strongly correlated with the fraction of planktonic cells it contains. In addition to revealing the existence of previously undocumented interactions within a vertebrate intestine, this exploratory experiment underscores how tools for genetically manipulating diverse bacterial symbionts facilitates comparative studies involving multiple species.

In total, the tools and step by step protocols described here will empower a wide range of researchers studying different host-microbe systems as well as free-living bacteria to explore deeper into the inner workings of wild and novel bacterial isolates. Our solutions equally enhance the genetic manipulation of both established and newly emerging model bacterial lineages and will speed the research pipeline from metagenomics to mechanistic microbiology.

## RESULTS

### The lack of compatible post-conjugation counterselection strategies and the deleterious nature of bacterial domestication

The process of genetically manipulating a bacterium typically begins with the delivery of recombinant DNA into its cell. Several methods can be used for this purpose, but conjugation or bacterial mating—which is the transfer of DNA from a donor cell to a target cell—is highly efficient and versatile, working with a wide range of bacterial lineages. During this procedure, successfully modified target cells are made drug resistant to facilitate their selective recovery. However, a constraint of conjugation is that it also depends on a strategy for simultaneously counterselecting against the donor cells that harbor the same drug resistance. This can be problematic when dealing with novel and uncharacterized bacteria because counterselection schemes often rely on known aspects of a target strain’s physiology. For example, many donor strains (e.g., *E. coli* SM10) are auxotrophic for amino acids and vitamins and thus, can be counterselected on defined growth media lacking specific metabolites. But for this approach to work the target strain cannot itself be auxotrophic and the right mixture of nutrients and ions must be formulated, which is not always straightforward due to the complex metabolic needs of different bacteria. We found that this drawback represents an immediate hurdle when working with several zebrafish-derived bacterial symbionts. None of the isolates we aimed to genetically manipulate could grow on a standardized defined growth medium (i.e., M9 minimal media) that is used for *E. coli* SM10 counterselection. This ultimately meant that customized counterselection media would need to be painstakingly developed for each new target strain.

Faced with this scenario, we turned to domestication, which is the process of modifying target strains with a selective trait that is not expressed by the donor. This is commonly done by isolating target strain variants with spontaneous antibiotic resistance. However, despite the wide use of this technique, antibiotic resistances associated with domestication (e.g., to rifampicin or streptomycin) typically arise because of mutations within critical cellular machinery, such as RNA polymerase or components of the ribosome, which can severely impair the natural physiology of bacteria and render them unsuitable for study^19–25^. Consistent with the deleterious consequences of domestication, we found that a rifampicin-resistant variant of the zebrafish intestinal symbiont *Vibrio cholerae* ZWU0020, denoted ZWU0020R, exhibits highly perturbed growth kinetics in vitro (Figure 1A). In addition, although domestication of *Vibrio* ZWU0020 did allow us to successfully insert a gene encoding green fluorescent protein (GFP) within its genome, this modification further aggravated its poor growth phenotype (Figure 1A). As another assessment of ZWU0020R’s altered physiology, we characterized its motility phenotype and observed that it displays attenuated swimming in soft agar compared to wild-type (Figure 1B). To determine if altered swimming is a phenotype of all rifampicin-resistant *Vibrio* ZWU0020 variants, we inspected the motility phenotype of four independently derived clones. Interestingly, two of the clones are attenuated like the original ZWU0020R strain, whereas the other two perform similar to wild-type, suggesting that they may carry alternative and/or compensatory mutations (Figure 1C). Altogether, our experience attempting to genetically manipulate a variety of novel zebrafish bacterial symbionts using conventional approaches highlights the limitations of current counterselection strategies and the deleterious nature of domestication.

**Figure 1.**
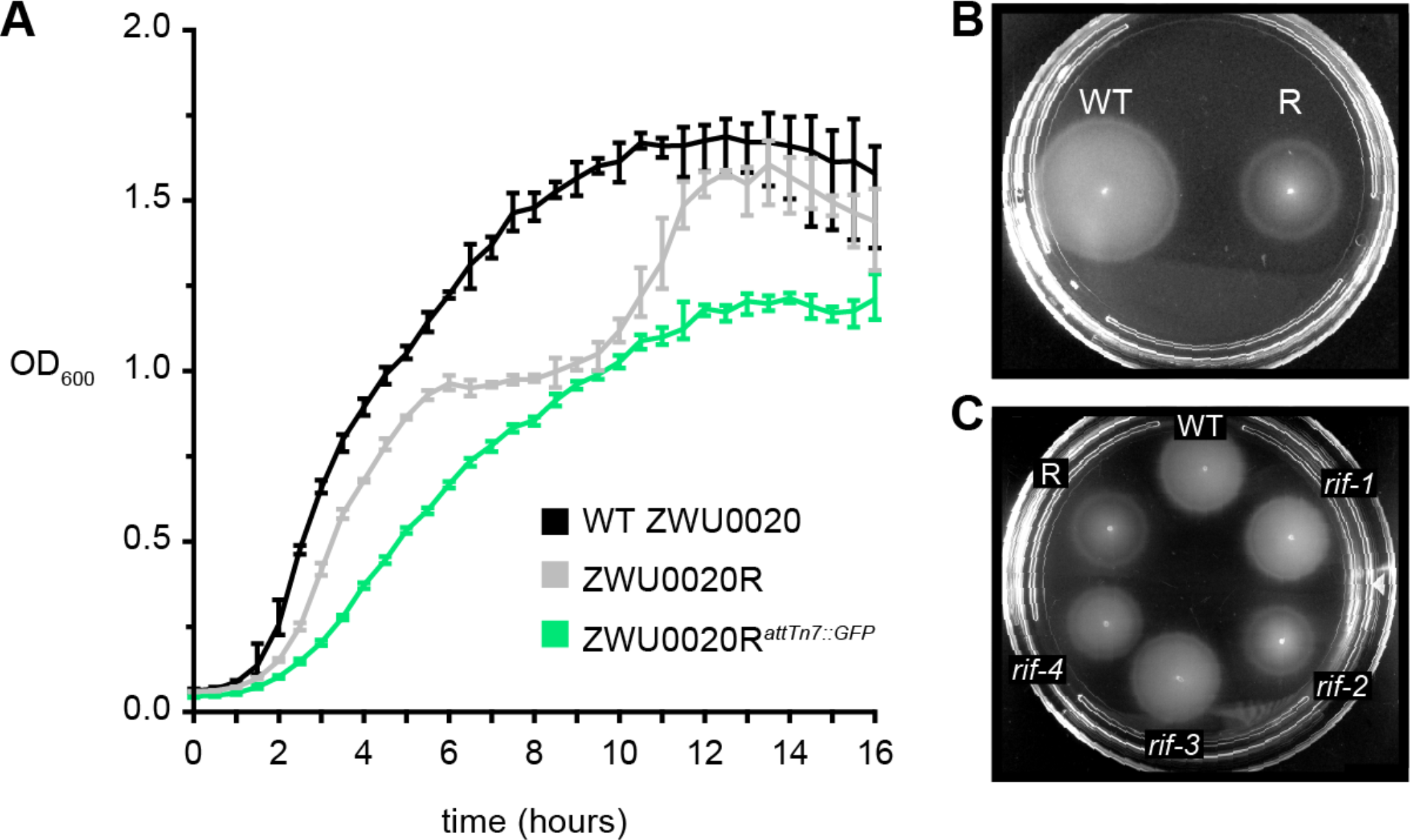
Domestication results in physiological defects. (**A**) Plotted is the average optical density at 600nm (OD_600_) vs. time (hours) of wild-type (WT) *Vibrio* ZWU0020, and its rifampicin-domesticated derivatives ZWU0020R and ZWU0020R^*attTn7*::*GFP*^, during shaking growth in LB broth at 30°C. Range bars are based on four technical replicates. (**B**) Swim motility of WT and ZWU0020R (R) in 0.2% tryptic soy agar at 30°C. (**C**) Swim motility of four rifampicin-resistant (*rif*) *Vibrio* ZWU0020 variants compared to WT and ZWU0020R performed as in **B**.

### Temperature and kill switch-based systems for domestication-free counterselection of donor cells

To address the lack of adequate post-conjugation counterselection methods, we set out to develop strategies that are technically straightforward and not reliant on inherent or domesticated traits of target strains. We devised two plasmid-based counterselection systems that control donor cell growth by a mechanism similar to that of common suicide vectors. The first system is temperature-based and works through a temperature-sensitive origin of replication that restricts donor cell growth in the presence of antibiotic selection at or above 37°C. Temperature-based control of plasmid replication is well established, but has not been widely implemented as a method of post-conjugation counterselection despite its amenability and previous indications that it can be used in this way^26^. The second system restricts donor cell growth through a genetic kill switch that, when induced, leads to the expression of three toxic peptides. These two approaches differ in their mode of action and offer slightly different procedural advantages. Notably, we chose to develop plasmid-based counterselection systems because their portability allows the use of alternative donor strains. To initially test the utility of each counterselection system, we incorporated them into existing vectors that are commonly used for making targeted Tn7 transposon-based chromosomal insertions^27^.

Temperature-based counterselection was achieved by replacing the R6K origin of replication of the Tn7-tagging vector pUC18R6KT-mini-Tn7T-GM (pTW56) with the temperature-sensitive origin of replication *ori*_*101*_/*repA*101^ts 28^ (Figure 2A). The resulting vector, pTn7xTS (**<u>T</u>**emperature-**<u>S</u>**ensitive), mediates temperature-dependent growth of *E. coli* SM10 (Figure 2-Figure Supplement 1). At the permissive temperature of 30°C, SM10/pTn7xTS grows normally on rich media in the presence of antibiotic selection. At the restrictive temperature of 37°C, the vector is unable to be maintained, leading to loss of antibiotic resistance and a drop in viability by up to three orders of magnitude (Figure 2-Figure Supplement 1). In the context of an example Tn7-tagging protocol, conjugation is performed at 30°C without antibiotic selection between two SM10 donor strains and a *Vibrio* target strain (Figure 2B, left). The SM10 donors carry either pTn7xTS (donor^Tn^) or the transposase-encoding helper plasmid pTNS2 (donor^helper^). At this point in the procedure, only the donor^Tn^ strain is resistant to the selective antibiotic being used, which in this scenario is gentamicin. Successfully modified *Vibrio* cells harboring a chromosomal copy of the Tn7 transposon, along with the gentamicin resistance gene it encodes, are then selected for by plating the mating mixture in the presence of gentamicin at 37°C (Figure 2B, right). The donor^Tn^ strain is counterselected because it is unable to maintain plasmid-based resistance at 37°C, whereas the donor^helper^ strain remains sensitive to gentamicin throughout the procedure.

A strength of temperature-based counterselection is that it is technically simple, requiring only a shift in growth temperature, but it is limited to target strains that can grow at 37°C. This constraint is problematic for several bacterial lineages native to zebrafish as well as other ectotherms, such as stickleback or fruit flies, which cannot survive at temperatures above the growth temperature of their host (in these cases, ≤ 30°C). Therefore, we developed a second strategy based on an inducible kill switch that functions independently of growth temperature.

The kill switch we designed consists of two elements: a constitutively expressed *lacl* gene, which encodes the lac repressor, and a synthetic operon containing three *E. coli*-derived genes encoding toxic peptides—HokB, GhoT, and TisB—placed under the control of the Lacl-repressible promoter P_tac_ (Figure 2-Figure Supplement 2A)^29–33^. Upon induction by the allolactose analogue isopropyl-β-D-thiogalactoside (IPTG) these toxic peptides act to disrupt the proton-motive force within donor cells, leading to impaired ATP synthesis and death. We built this kill switch counterselection system into the backbone of the Tn7-tagging vector pUC18T-mini-Tn7T-GM (pTW54), producing pTn7xKS (**<u>K</u>**ill **<u>S</u>**witch) (Figure 2C). In the presence of antibiotic selection and IPTG, pTn7xKS is capable of inhibiting SM10 growth by up to four orders of magnitude (Figure 2-Figure Supplement 2B). Of note, initial kill switch prototypes carrying only a single toxin gene were less potent, which may be an important consideration in future kill switch designs (Figure 2-Figure Supplement 2B). In the context of a Tn7-tagging scenario, kill switch-based counterselection is carried out in much the same way as temperature-based counterselection, except that selection of modified target cells is done on media containing IPTG at a growth temperature suitable for the target strain being used (Figure 2D).

**Figure 2.**
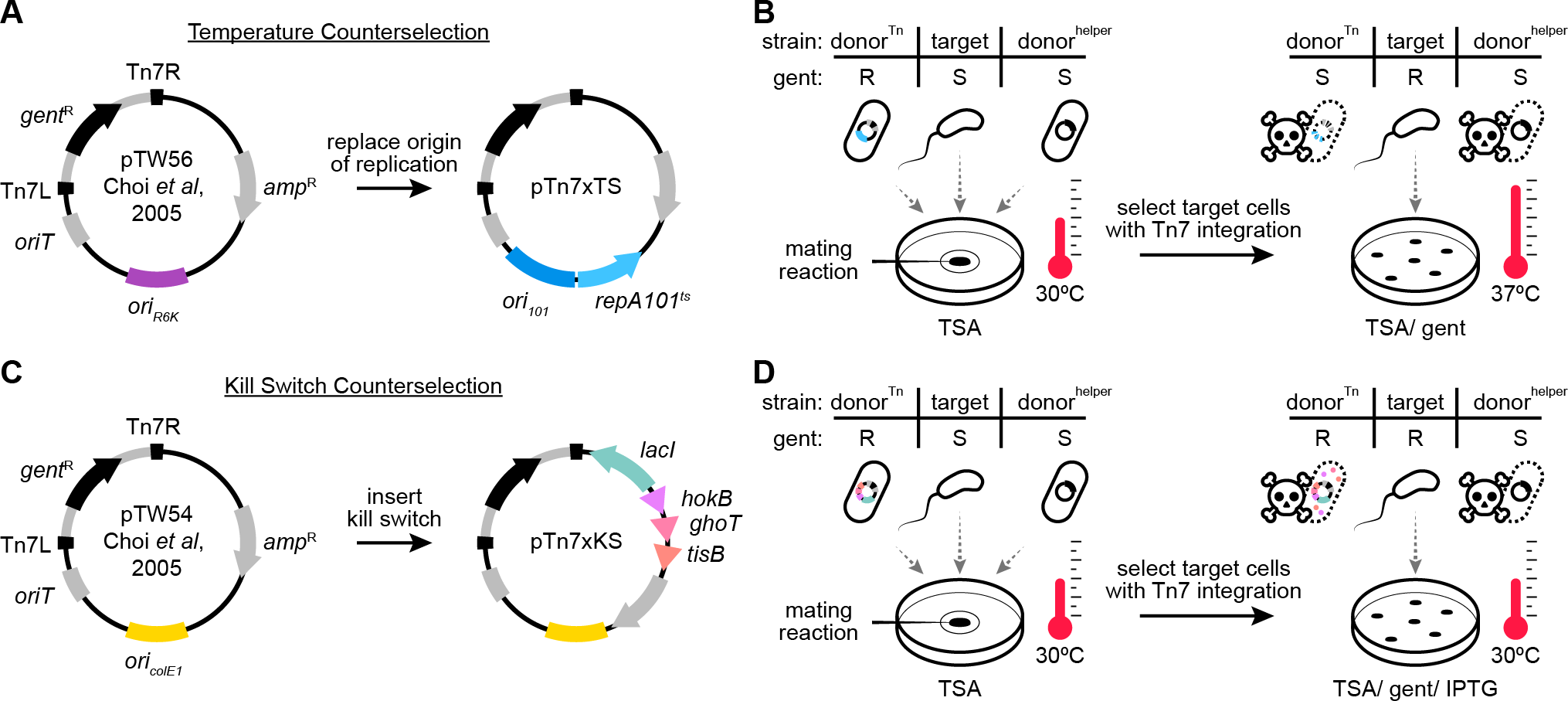
Construction and application of domestication-free counterselection systems. (**A**) Temperature-based counterselection was achieved by replacing the R6K origin of replication (*ori*_*R6K*_) of pUC18R6KT-mini-Tn7T-GM (pTW56) with the temperature-sensitive origin of replication *ori*_*101*_/*repA*101^ts^. Tn7L and Tn7R inverted repeats flank the Tn7 transposon (gray stroke). *gent*^R^, gentamicin resistance gene; *amp*^R^, ampicillin resistance gene; *oriT*, origin of transfer. (**B**) Left: triparental conjugation between SM10 donor strains carrying either a temperature-sensitive Tn7-tagging vector (donor^Tn^) or transposase helper vector (donor^helper^) and a *Vibrio* target strain. Gentamicin (gent) phenotype of each strain is indicated as resistant (R) or sensitive (S). Mating reactions are incubated at 30°C upon a filter disc on a trypic soy agar (TSA) plate. Right: post-conjugation counterselection of donor cells is done on TSA/ gent plates at 37°C. (**C**) Kill switch-based counterselection was achieved by inserting a Lacl-regulated toxin array, comprised of the genes *hokB*, *ghoT*, and *tisB*, into the backbone of pUC18T-mini-Tn7T-GM (pTW54). *ori*_*ColE1*_, high copy number origin of replication. (**D**) Left: triparental conjugation as in **B**, except donor^Tn^ carries a kill switch Tn7-tagging vector. Right: post-conjugation counterselection of donor cells is done on TSA/ gent/ IPTG plates at 30°C.

### Chromosomal insertion of rationally designed gene expression scaffolds into wild and uncharacterized bacterial lineages using domestication-free counterselection systems

To test the effectiveness of our domestication-free counterselection systems, we employed them to integrate genetically encoded fluorescent proteins into the chromosome of various uncharacterized zebrafish bacterial symbionts. However, while exploring available gene expression constructs, we found that many are inflexible and inadequately designed. Specifically, vectors often contain extraneous DNA sequences left over from previous imprecise subcloning procedures and have little to no options for customizing important sequence motifs. The ability to customize expression constructs is critical when working with lineages that differ in, for instance, optimal promoter sequences or ribosome binding sites. Therefore, we first addressed the need for standardized expression constructs by rationally designing a modular gene expression scaffold.

An expression scaffold containing four interchangeable elements—a promoter, 5’ and 3’ untranslated regions (UTR), and an open reading frame (ORF)—was built into the multiple cloning site (mcs) of pGEN-mcs^34^, producing pXS (e**<u>X</u>**pression **<u>S</u>**caffold) (Figure 3). pGEN-mcs was chosen to house the expression scaffold because it enables fast and easy prototyping of scaffold parts in *E. coli*, which like many zebrafish bacterial symbionts, is a member of the Gammaproteobacteria and shares basic genetic control elements. Restriction sites underlie the modular architecture of the scaffold and allow each part to be customized (Figure 3-Figure Supplement 1A). Sequence motifs can be replaced individually or the entire scaffold can be subcloned. As initially built, a minimal P_tac_ promoter without the lac operator sequence, which avoids interference from an endogenously encoded lac repressor if present, is used to achieve constitutive transcription. A synthetic 5’ UTR containing both an epsilon enhancer sequence and ribosome binding site controls translation^35,36^. The 3’ UTR, which was originally present within pGEN-mcs, contains a *trpL* attenuator sequence for transcriptional termination^37^. Lastly, three different ORFs encoding the fluorescent proteins sfGFP^38^, dTomato^39^, and mPlum^40^ were each used to produce three separate expression scaffold variants, pXS-sfGFP, pXS-dTomato, and pXS-mPlum (Figure 3-Figure Supplement 1A). After assembly, each expression scaffold was subcloned into Tn7-tagging vectors with either temperature (i.e., pTn7xTS) or kill switch (i.e., pTn7xKS) counterselection systems (Figure 3-Figure Supplement 1B and C).

Tn7-tagging vectors equipped with rationally designed expression scaffolds were next used to carry out chromosomal tagging of *Vibrio* ZWU0020 as outlined in Figure 2B and 2D. Unlike previous attempts (Figure 1A), domestication-free manipulation of *Vibrio* ZWU0020—using either temperature or kill switch counterselection systems—both worked and preserved this strain’s normal physiology (Figure 3-Figure Supplement 2). To demonstrate the compatibility of these tools across different zebrafish symbiont lineages, we tagged 10 strains representing 6 different genera—including *Vibrio*, *Aeromonas*, *Pseudomonas*, *Acinetobacter*, *Enterobacter*, and *Plesiomonas* (Supplementary File 1). Multiple variants that express either *sfGFP*, *dTomato*, or *mPlum* were generated for many of the lineages (Supplementary File 1). This collection of marked zebrafish symbionts serves as a resource of ready-made strains for the zebrafish microbiota community. Notably, several of the lineages manipulated are novel and have yet to be assigned species designations, which is typical of symbionts associated with complex host-associated communities. We also verified that domestication-free counterselection systems can be used to manipulate other species not associated with the zebrafish intestinal microbiota, including a human-derived strain of *E. coli* (HS)^41^, the leech symbiont *Aeromonas veronii* HM21^42^, and an environmental isolate of *Shewanella oneidensis* (MR-1)^43^ (Supplementary File 1).

**Figure 3.**
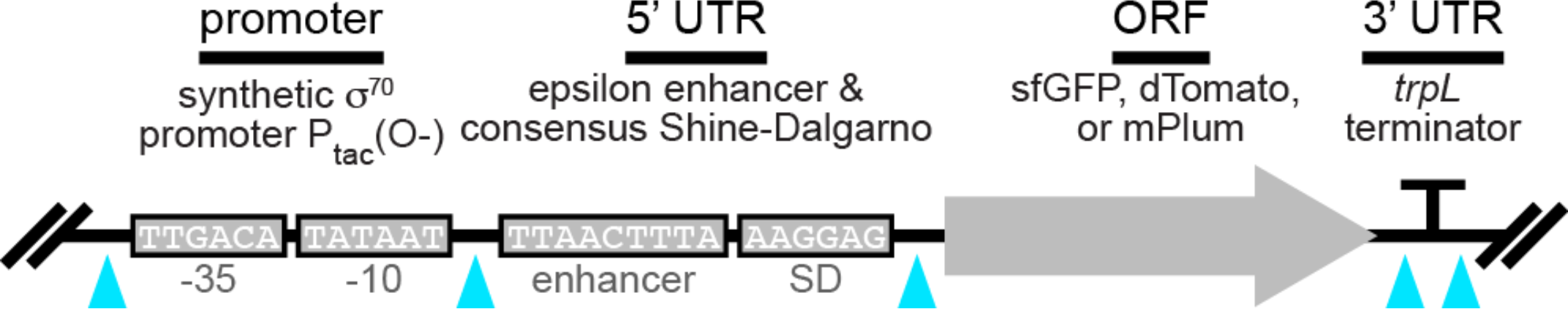
Gene expression scaffold design features. Each interchangeable element is flanked by restriction sites (cyan arrowheads). Promoter: constitutively active P_tac_ promoter without lac operator sequence (O-) drives transcription. 5’ untranslated region (UTR): epsilon enhancer sequence and consensus ribosome binding site (i.e., Shine-Dalgarno sequence) promote strong translation. Open reading frame (ORF): encodes a single fluorescent protein. 3’ UTR: *trpL* attenuator sequence terminates transcription.

### Streamlining allelic exchange by visualizing homologous recombination events using a fluorescent tracker

Allelic exchange is a robust and versatile homologous recombination technique for making targeted genetic knock-ins and knock-outs in bacteria^44–46^. Therefore, to extend the utility of our domestication-free counterselection systems, we incorporated them into currently available vectors that are used for mediating allelic exchange. As expected, these updates facilitated the domestication-free engineering of gene deletions in several uncharacterized symbiotic bacteria. However, not all bacteria tested could be successfully manipulated using current allelic exchange protocols, highlighting another breakdown in the compatibility.

Allelic exchange involves two successive homologous recombination events between an allelic exchange vector and the bacterial chromosome. The crux of allelic exchange is isolating rare unmarked mutant cells from large populations of heterozygous intermediates known as merodiploids that arise after the vector integrates into the chromosome during the first recombination step. A longstanding strategy for recovering variants that have undergone the second recombination, which results in vector loss, works by restricting merodiploid growth. This is typically done by expressing a gene called *sacB* located within the allelic exchange vector backbone that confers growth inhibition in the presence of sucrose^47^. Although widely used, *sacB* counterselection of merodiploids does not always work and can be difficult to troubleshoot when it fails. We experienced these shortcomings while attempting to delete a gene associated with chemotactic behavior in a zebrafish symbiont, *Vibrio furnissii* ZOR0035, using the common *sacB*-based allelic exchange vector pDMS197^48^. *Vibrio* ZOR0035 merodiploids are refractory to *sacB* counterselection, which made it impossible to isolate cells with the desired mutation. We surmise that the counterselection fails in some bacterial lineages because the expression or activity of the levansucrase enzyme encoded by *sacB*, which synthesizes high-molecular-weight fructose polymers, is inadequate. To overcome lineage-specific limitations of *sacB* counterselection, we developed a more tractable strategy based on visual markers.

Our solution uses GFP to track the merodiploid status of target cells (Figure 4A). In this way, the initial recombination step generates GFP-positive merodiploid populations that can be readily screened for cells where the second recombination step has occurred, producing GFP-negative mutants (i.e., instances of “successful” allelic exchange), which typically occur with equal frequency as wild-type revertants (i.e., instances of “aborted” allelic exchange) (Figure 4A). To test the feasibility of this approach, we revisited the engineering of a gene deletion in *Vibrio* ZOR0035. A constitutively expressed GFP gene was inserted into the backbone of a prototype pDMS197 vector containing a kill switch counterselection system and an allelic exchange cassette targeting the chemotaxis gene *cheA*. At the time of this work, *cheA* was the focus of an unrelated project, and it is used here merely to demonstrate proof of concept. As outlined in Figure 4B, the GFP marked allelic exchange vector was delivered into *Vibrio* ZOR0035 via conjugation. GFP-positive merodiploids, harboring an integrated copy of the allelic exchange vector, were readily isolated and purified. Of note, over the course of this work we empirically determined that the kill switch toxins do not interfere with merodiploid growth in several different bacteria, indicating that either they have restricted activity and are only lethal to *E. coli* donor cells or they fail to reach toxic levels when expressed from a single chromosomal locus. Next, populations of merodiploids were expanded in liquid culture and plated on nonselective media at a density that allowed discrete colonies to form. Colonies exhibiting sectored regions of GFP loss were then purified to obtain isogenic clones, and putative mutants were genotyped by polymerase chain reaction (PCR). Genotyping was done using PCR primers that flank the *cheA* locus, yielding a single large amplification product if the *cheA* gene is present and a smaller sized product if the mutant allele is present. Because they are heterozygous, merodiploids produce both products. Ultimately, our visual merodiploid tracking strategy proved extremely efficient and straightforward to perform, allowing us to successfully engineer a targeted gene deletion in a bacterial strain that was otherwise genetically intractable using previous methodologies.

**Figure 4.**
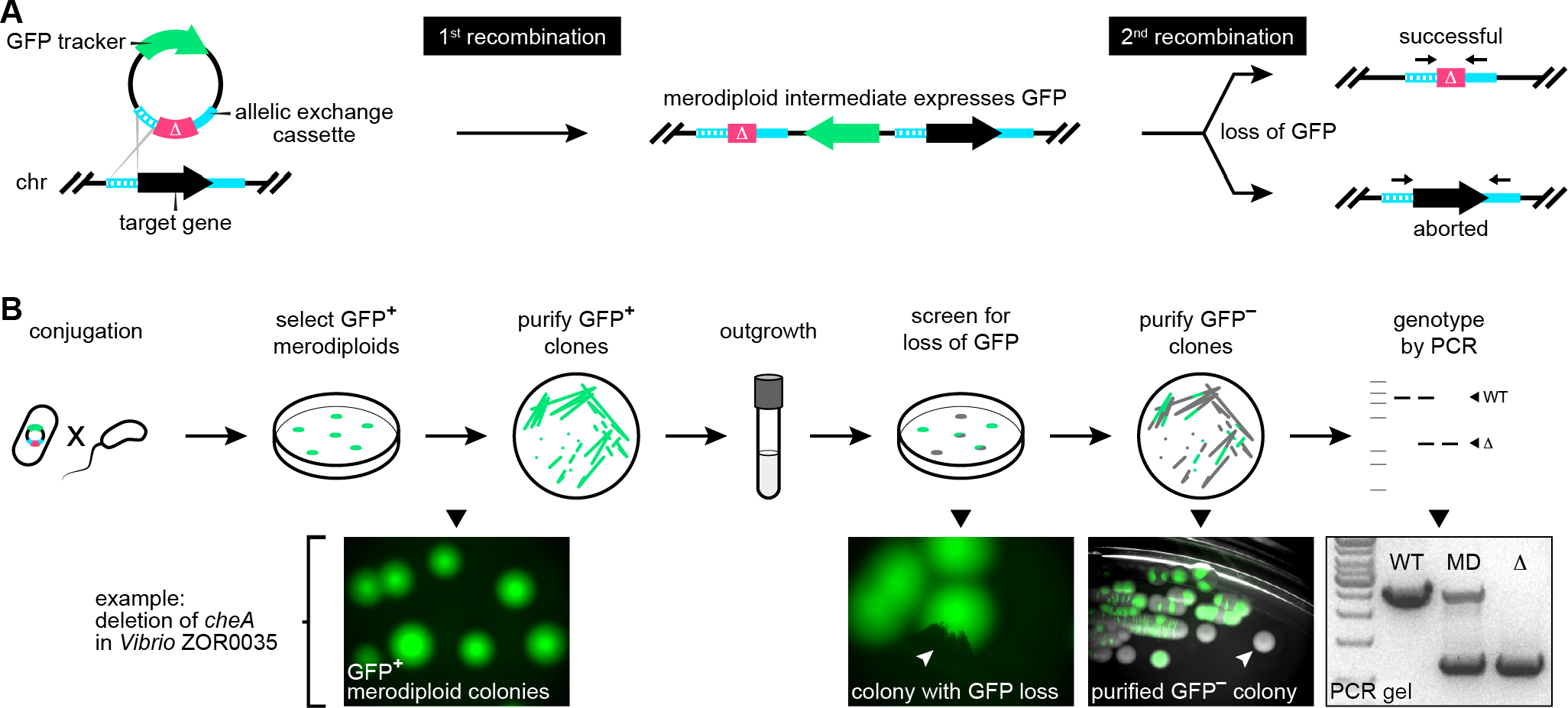
Performing allelic exchange with a fluorescent merodiploid tracker. (**A**) Outline of recombination events during allelic exchange with a fluorescent tracker. Depicted is an allelic exchange vector that expresses GFP and carries a cassette comprised of a mutant allele (Δ, magenta) flanked by regions (hashed and solid cyan strokes) homologous to regions flanking a target gene located on the bacterial chromosome (chr). The first recombination event—which randomly occurs between either homology region—integrates the vector into the chromosome, producing a GFP-expressing merodiploid. The second recombination event results in GFP loss. If it occurs between the unused homology region (i.e., the “solid” region in this scenario), then allelic exchange is successful. If it occurs between the same region (i.e., the “hashed” region), the original wild-type locus is restored. Black arrows above final allelic exchange products denote primer annealing sites for PCR-based genotyping depicted in **B**. (**B**) Top row illustrates the procedural steps of allelic exchange using a fluorescent merodiploid tracker. Bottom row shows example images acquired during the engineering of a gene deletion in *Vibrio* ZOR0035. White arrowheads indicate colonies with partial or complete loss of GFP expression. WT, wild-type *Vibrio* ZOR0035; MD, merodiploid; Δ, Δ*cheA* mutant.

### Gene deletion and complementation with modernized engineering vectors

To complete the genetic toolkit for manipulating wild and diverse bacterial isolates, we combined the tools and approaches described thus far to construct a set of adaptable allelic exchange vectors that further improve the tractability of making markerless genetic alterations. These modernized vectors incorporate fluorescent merodiploid tracking and domestication-free counterselection systems within a highly customizable plasmid architecture (Figure 5, “vector design”). Molecular scaffolds for holding antibiotic selection markers, fluorescent trackers, and a counterselection kill switch were built into a pUC-derived vector backbone that has a temperature-sensitive origin of replication and a single blunt restriction site for straightforward insertion of allelic exchange cassettes. This modular design allows virtually every functional element to be customized for different bacterial lineages (Figure 5-Figure Supplement 1).

In total, two allelic exchange vectors were generated, pAX1 and pAX2 (**<u>A</u>**llelic e**<u>X</u>**change), which differ in their domestication-free counterselection systems (Figure 5). Both vectors mediate temperature-based counterselection of SM10 donor cells, but pAX2 also contains a TetR-regulated kill switch that can be induced by anhydrotetracycline (Figure 5-Figure Supplement 2). Notably, the dual temperature/kill switch counterselection activity of pAX2 is quite potent, reducing SM10 viability by over five orders of magnitude (Figure 5-Figure Supplement 2B). Two resistance markers for gentamicin and chloramphenicol were included to give pAX1 and pAX2 greater “off the shelf” compatibility with different target strains.

**Figure 5.**
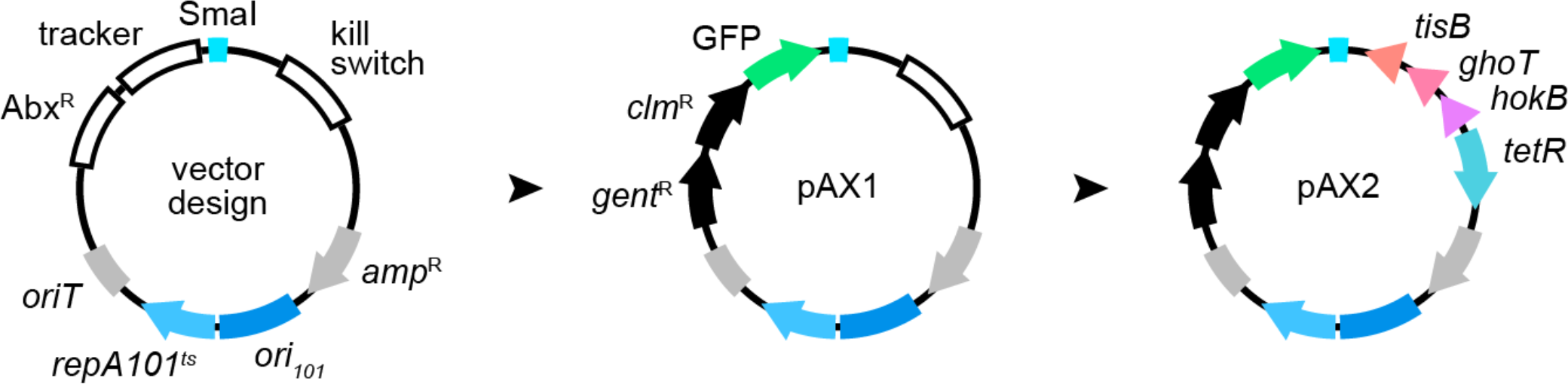
Rational design of customizable allelic exchange vectors. “vector design” illustrates vector architecture. Features include customizable molecular scaffolds for holding antibiotic selection markers (Abx^R^), a merodiploid tracker, a single Smal restriction site for insertion of allelic exchange cassettes, and an option for kill switch-based counterselection of donor cells. pAX1 was initially constructed, and carries two antibiotic selection markers encoding resistance to gentamicin (*gent*^R^) and chloramphenicol (*clm*^R^) along with a GFP tracker. pAX2 was derived via the insertion of a tet-inducible kill switch. *oriT*, origin of transfer; *ori*_*101*_/*repA*101^ts^, temperature-sensitive origin of replication; *amp*^R^, ampicillin resistance gene.

These new allelic exchange vectors were next used to engineer markerless gene deletions. For this proof of concept, we designed an allelic exchange cassette to delete two neighboring genes in *Vibrio* ZWU0020, *pomA* and *pomB*, which encode components of the polar flagellar motor. After inserting the cassette into an early, but functionally equivalent, version of pAX1 (see Materials and Methods), GFP-positive merodiploids were generated and isolated as before and screened for loss of GFP expression and thus, the integrated vector (Figure 6-Figure Supplement 1). Mutants harboring the desired mutation, which fused the start codon of *pomA* with the stop codon of *pomB*, were confirmed by PCR (Figure 6A and 6B). As anticipated, ZWU0020 Δ*pomAB* exhibited complete loss of swimming motility in soft agar (Figure 6C, bottom left) without overt growth defects in liquid media (Figure 6-Figure Supplement 1). To demonstrate the cross-lineage compatibility of these tools, we successfully employed pAX2 to create a similar markerless deletion of a homologous *pomAB* locus in another zebrafish symbiont, *Aeromonas veronii* ZOR0001 (Figure 6-Figure Supplement 2). Notably, while screening *Vibrio* ZWU0020 and *Aeromonas* ZOR0001 merodiploid colonies for putative mutants, we observed that many different patterns of GFP loss can arise (Figure 6- Figure Supplement 3). Remarkably, even in situations where mutant cells reside within miniscule GFP-negative patches, a single colony purification step can often be used to isolate them. The ability to readily identify and recover mutant cells with such sensitivity highlights the robustness of our visual screening approach.

Constructing deletion mutants is just the first step in dissecting the genetic pathways that underlie a given activity or behavior. Complementation must also be performed to rigorously confirm that a mutant phenotype is the result of a specific genetic disruption and not polar effects or other unintended consequences of chromosomal manipulation^15^. Therefore, we wanted to demonstrate how our domestication-free Tn7-tagging vectors could be employed to complement the ZWU0020 Δ*pomAB* mutant. The *pomAB* locus of ZWU0020, including the native *pomA* promoter, was PCR-amplified and inserted within the Tn7 transposon of the tagging vector pTn7xTS-sfGFP, which also contains a constitutively expressed *sfGFP* (Figure 6D). Chromosomal insertion of this construct at the *attTn7* site of ZWU0020 Δ*pomAB* fully restores wild-type motility, thus confirming that sole disruption of these genes caused the loss of motility in the mutant (Figure 6C, bottom right).

**Figure 6.**
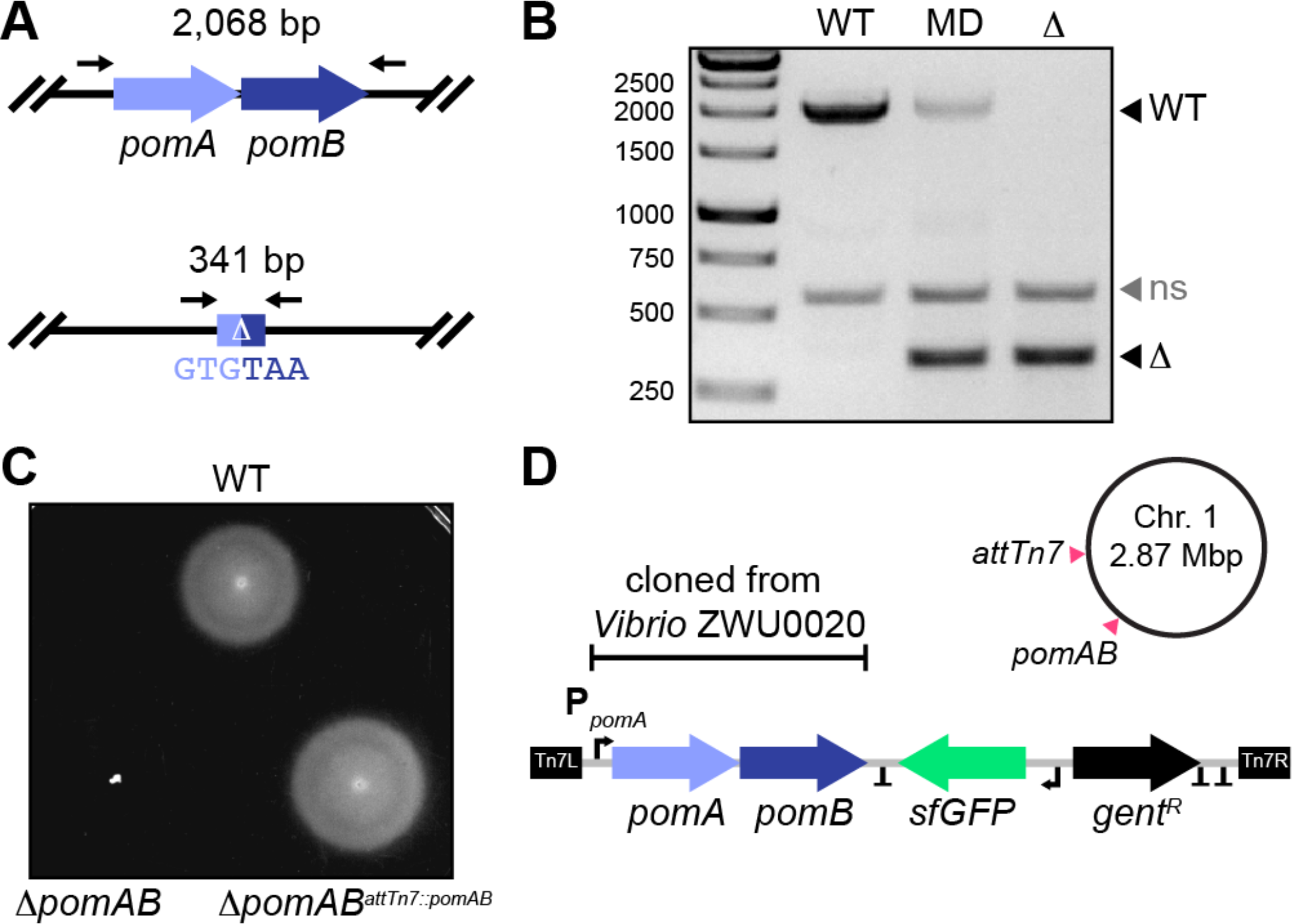
Gene deletion and complementation with modernized engineering vectors. (**A**) Top: wild-type *pomAB* locus in *Vibrio* ZWU0020. Bottom: result of markerless *pomAB* deletion via allelic exchange. Black arrows mark approximate primer annealing sites for genotyping and the size of each amplification product is indicated. (**B**) Agarose gel showing PCR-based genotyping of wild-type (WT), merodiploid (MD), and a Δ*pomAB* (Δ) mutant. Migration distances of WT and mutant alleles are indicated. ns, nonspecific amplification product. (**C**) Swim motility of WT, Δ*pomAB*, and the complemented Δ*pomAB*^*attTn7*::*pomAB*^ variant in 0.2% tryptic soy agar at 30°C. (**D**) Shown is a schematic of the Tn7 transposon from pTn7xTS-sfGFP used for complementation, which was modified to carry the native *pomAB* locus of *Vibrio* ZWU0020. Also depicted are the relative positions of where the *pomAB* genes were deleted and reintroduced at the Tn7 insertion site (*attTn7*) on chromosome 1 of Vibrio ZWU0020. “T” denotes transcriptional terminators; Tn7L and Tn7R, Tn7 inverted repeats, P_pomA_, native *pomA* promoter; *gent*^R^, gentamicin resistance gene; sfGFP, fluorescent tag.

### Live imaging of diverse bacterial symbionts yields insights into host-microbe interactions within the vertebrate intestine

The power to genetically manipulate a diverse range of bacterial symbionts provides an opportunity for comparative studies focused on identifying unique or broadly conserved features of host–microbe interactions. To illustrate how the tools developed in this work facilitate such investigations, we examined the intestinal colonization patterns and cellular behaviors of seven fluorescently tagged zebrafish symbionts by light sheet fluorescence microscopy^13,49^. The strains chosen included: *Aeromonas* ZOR0001, *Aeromonas* ZOR0002, *Enterobacter* ZOR0014, *Plesiomonas* ZOR0011, *Pseudomonas* ZWU0006, *Vibrio* ZOR0036, and *Vibrio* ZWU0020. A tagged version of the ZWU0020 Δ*pomAB* motility mutant was also analyzed. Prior to imaging, bacteria were associated individually with 4-day old, germ-free larval zebrafish for 24 hours. Mono-association provides unrestricted access to the intestinal environment free of competition with resident microbiota, and therefore allows interactions between a single symbiont and its host to be studied in isolation. For each strain, real-time two-dimensional movies and three-dimensional images spanning the entire volume of the larval intestine were acquired from three separate hosts (Figure 7, Figure 7-Figure Supplement 1, and Supplementary Movies 1-8). From these data, distinct population structures are readily identified, and can be summarized by three properties: cell motility, growth mode (i.e., planktonic vs aggregated), and biogeography. Additional features of each strain, including estimated abundance, are provided in Figure 7-Figure Supplement 2.

**Figure 7.**
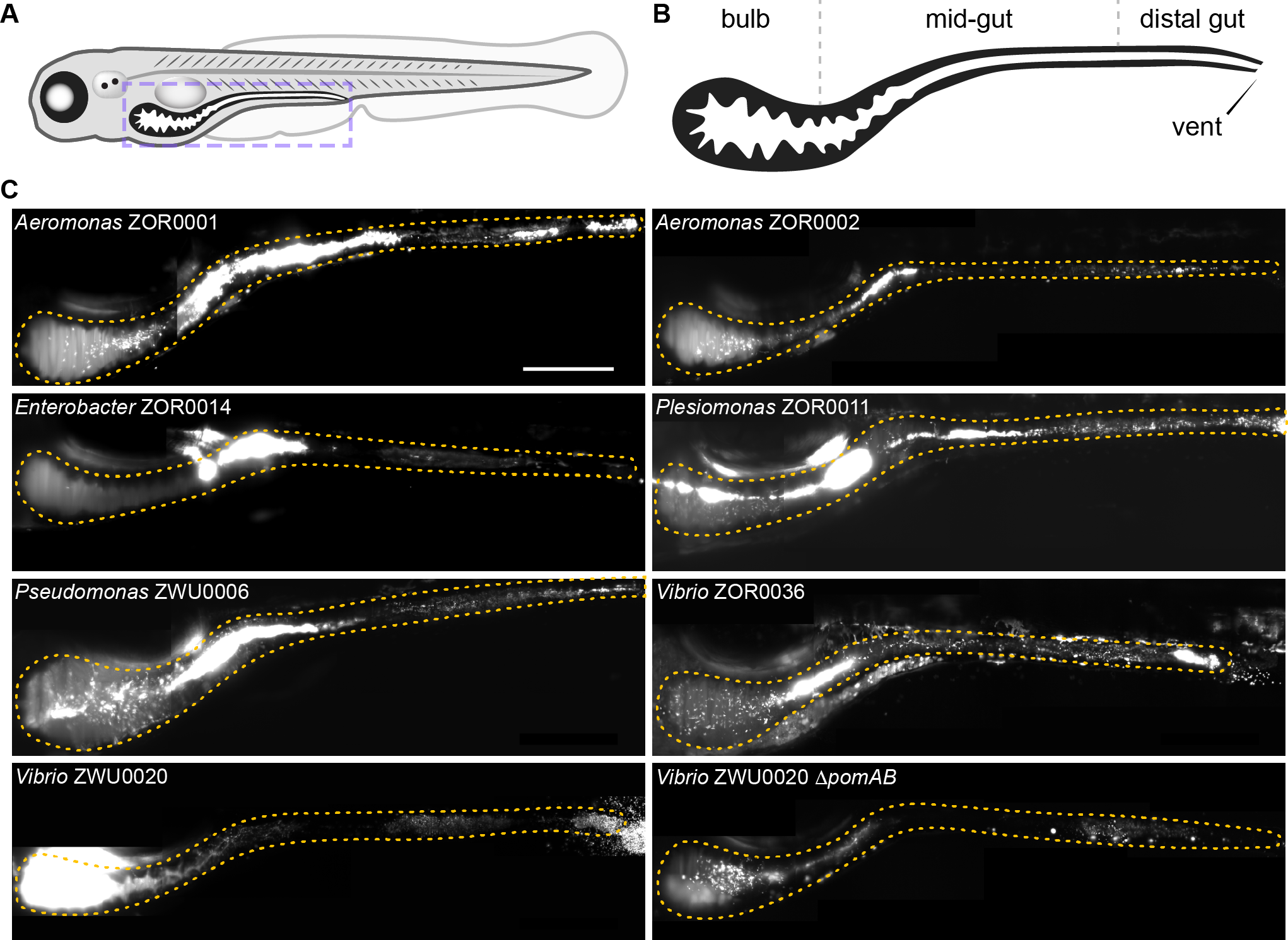
Intestinal colonization patterns and growth modes of zebrafish symbionts. (**A**) Cartoon diagram of a 5-day old larval zebrafish. Purple dashed box outlines region imaged in **C**. (**B**) Diagram shows the boundaries of the bulb, mid-gut, and distal gut within the larval intestine. The estimated bulb to mid-gut boundary is located where the bulb begins to become patently narrow. The mid-gut to distal gut boundary is approximately located where intestinal epithelial cells begin transitioning to a more colonic epithelial cell type^50^. (**C**) Maximum intensity projections of 3D image stacks acquired by light sheet fluorescence microscopy for indicated bacterial strains. Orange dotted outline marks the intestine in each image. Scale bar: 200μm.

Four of the seven wild strains examined display active motility within the intestine. Surprisingly, we found several discrepancies between the motility phenotype of strains in vivo and their predicted capacity for motility based on phylogenetic relatedness, genome sequence, and performance during in vitro assays (Figure 7-Figure Supplement 3 and Supplementary Movies 1-7). Each strain carries genes for flagellum biogenesis and displays free-swimming motility in liquid media. In addition, all strains were found to swim in soft agar, with the exception of *Vibrio* ZOR0036 (Figure 7-Figure Supplement 4). Yet despite these attributes, motile individuals were not observed within intestinal populations of *Enterobacter* ZOR0014, *Aeromonas* ZOR0001, and *Aeromonas* ZOR0002 (Supplementary Movies 2-4). We note that in a previous live imaging study involving *Aeromonas* ZOR0001 motile cells were detected, but that their occurrence was incredibly rare^13^. In addition, although *Vibrio* ZOR0036 exhibits little motility in soft agar, it gives rise to a considerable number of highly motile cells in vivo (Supplementary Movie 5); however, its overall motility phenotype is muted compared to closely related *Vibrio* ZWU0020 (Figure 7-Figure Supplement 3 and 4, and Supplementary Movie 1).

The growth mode of cells is categorized as either planktonic, which includes both motile and non-motile individuals, or aggregated. Both modes are typically represented across different populations, but the ratio of cells in each mode tends to be strain-specific. For example, at one extreme, populations of *Vibrio* ZWU0020 are almost entirely comprised of highly motile planktonic cells; this is more apparent in movies than three-dimensional image scans because of this strain’s high population density and fast movement (Supplementary Movie 1). At the opposite extreme, *Enterobacter* ZOR0014 forms large multicellular aggregates without motile individuals (Figure 7C and Supplementary Movie 2). The remaining strains produce populations with intermediate mixtures of planktonic and aggregated cells (Figure 7C and Figure 7-Figure Supplement 1).

Regarding biogeography, we observed a range of strain-specific spatial distributions along the length of the intestine. We can coarsely classify the location of bacterial populations as primarily residing in one of two regions, the proximal gut (referred to as the “bulb”) or the mid-gut, which we approximate in Figure 7B based on previous studies of larval zebrafish intestinal development^50,51^. For most strains, the bulk of their population is distributed throughout the mid-gut (e.g., *Aeromonas* ZOR0001, *Aeromonas* ZOR0002, *Enterobacter* ZOR0014, *Plesiomonas* ZOR0011, *Pseudomonas* ZWU0006, and *Vibrio* ZOR0036) (Figure 7C). By contrast, populations of *Vibrio* ZWU0020 are located within the proximal portion of the bulb, consistent with previous findings (Figure 7C)^13^. Notably, compared to wild-type, the ZWU0020 Δ*pomAB* motility mutant exhibits a reduction in overall population size and a slight shift in distribution to an area between the bulb and mid-gut (Figure 7C), indicating that for this strain, motility controls abundance and biogeography.

To distill and quantify our observations, we devised scalar metrics of biogeography and growth mode using computational image analysis (Materials and Methods). For each population, we computed the center of mass along the anterior-posterior axis of the intestine to represent biogeography and enumerated the fraction of cells that exist as planktonic individuals to represent growth mode. Plotting these data against each other shows a striking and unanticipated relationship: the fraction of planktonic cells within a strain’s population is strongly correlated with its biogeography (Figure 8). We quantified this relationship by linear regression of log-transformed planktonic fraction and population center medians, excluding the ZWU0020 Δ*pomAB* motility mutant (Figure 8, dashed trend line; *r*^2^ = 0.6762). Remarkably, the ZWU0020 Δ*pomAB* motility mutant conforms to this trend (Figure 8). As expected, populations of ZWU0020 Δ*pomAB* contain no motile individuals, but unexpectedly they adopt a more aggregated state compared to wild-type ZWU0020 (Supplementary Movie 8). This change in growth mode and the concomitant change in biogeography moves the ZWU0020 motility mutant along the trend line.

**Figure 8.**
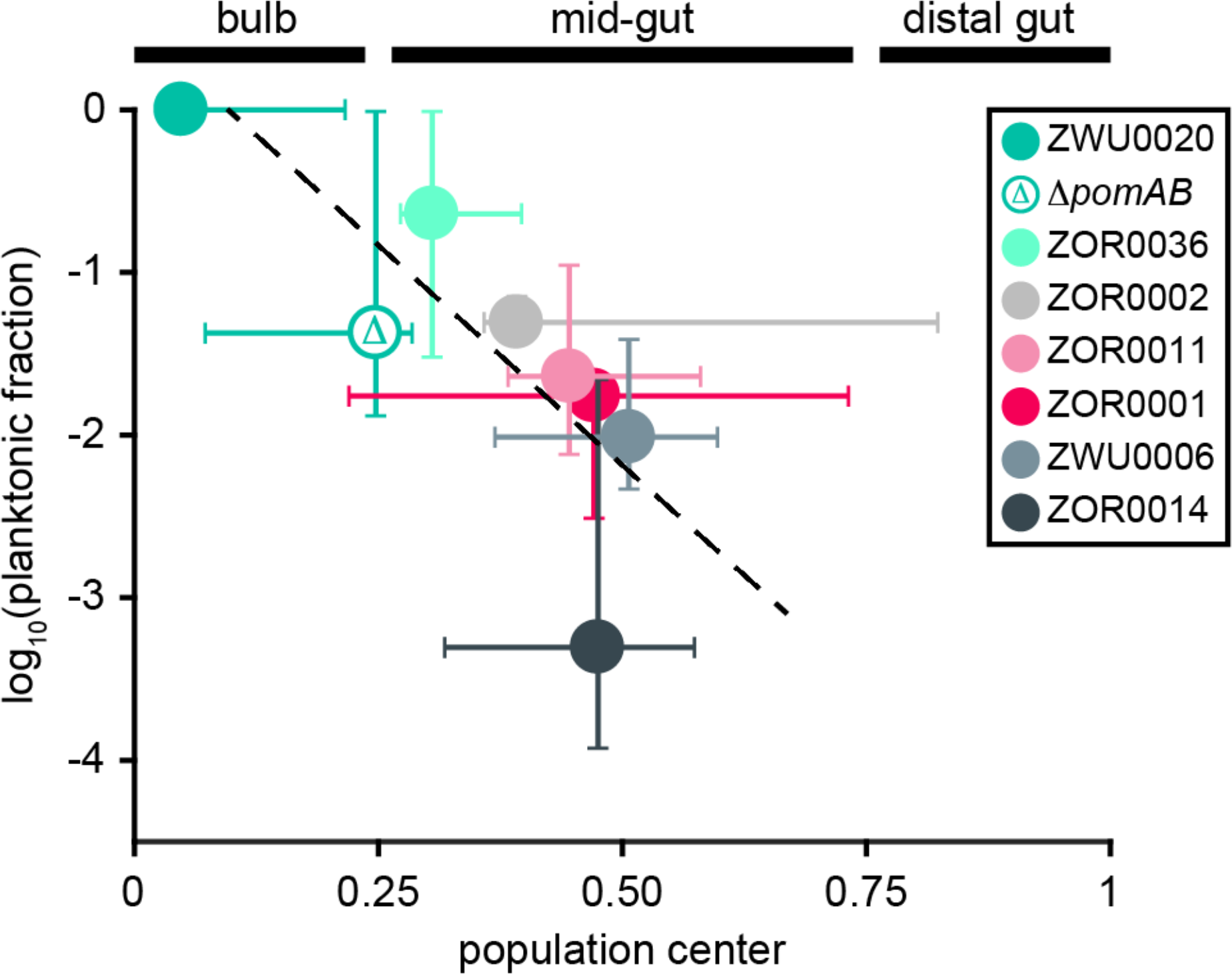
Relationship between growth mode and population biogeography. Plotted is the median log-transformed fraction of planktonic cells and median population 1D center of mass for each bacterial strain. Bars represent ranges. Data were derived from three animals (N = 3) per strain and are based on segmented 3D images from Figure 7 and Figure 7-Figure Supplement 1. Dashed trend line was generated by linear regression of median values (N = 7 data points, *r*^2^ = 0.6762). Corresponding boundaries for the bulb, mid-gut, and distal gut are indicated above the plot by black bars.

## DISCUSSION

### Impact on Bacterial Symbioses Research

Over the last decade, the landscape of bacterial symbiosis research has shifted due to an explosion in omics technologies and large-scale initiatives like the Human and Earth Microbiome Projects^52,53^. The traditionally static “one host, one microbe” view has given way to one that is more dynamic and complex, taking into account the highly contextual nature of symbiotic relationships and involvement of diverse multi-member microbial communities. This paradigm shift has generated several new challenges; chiefly, the demand for more efficient genetic manipulation of wild and novel symbiotic bacterial lineages. Addressing this problem is critical to experimentally unravelling the cellular and molecular determinants of host-microbe systems.

The impetus behind the genetic tools and approaches described in this work emerged from setbacks encountered while attempting to manipulate members of the zebrafish intestinal microbiota. The diversity of species and strains exposed several key inadequacies and weaknesses in conventional techniques, including the lack of domestication-free strategies for donor cell counterselection, poor modularity of available tools, and intractability of allelic exchange. Therefore, we designed tools and methods to circumvent these points of failure and quickly adapt to unforeseen idiosyncrasies of species or strains. In this way, an individual researcher or laboratory can focus on a single operating procedure using a centralized set of tools while being empowered to innovate when needed.

### Utility of the Tools Produced by this Work

An extensive collection of molecular tools is available for genetically manipulating bacteria; however, many can only be used with a small number of species or strains. The siloed nature of genetic tools puts a significant burden on researchers looking to manipulate diverse bacterial lineages because it forces them to sift through and become familiar with numerous different vectors and protocols. Furthermore, for those working with novel and uncharacterized bacteria or new to performing genetic manipulations altogether, developing a molecular starter kit is overwhelming. We addressed these problems by constructing a set of standardized engineering vectors that streamline the process of making genetic knock-ins and knock-outs across different lineages. These tools are briefly summarized in Figure 9 and their features are discussed below.

**Figure 9.**
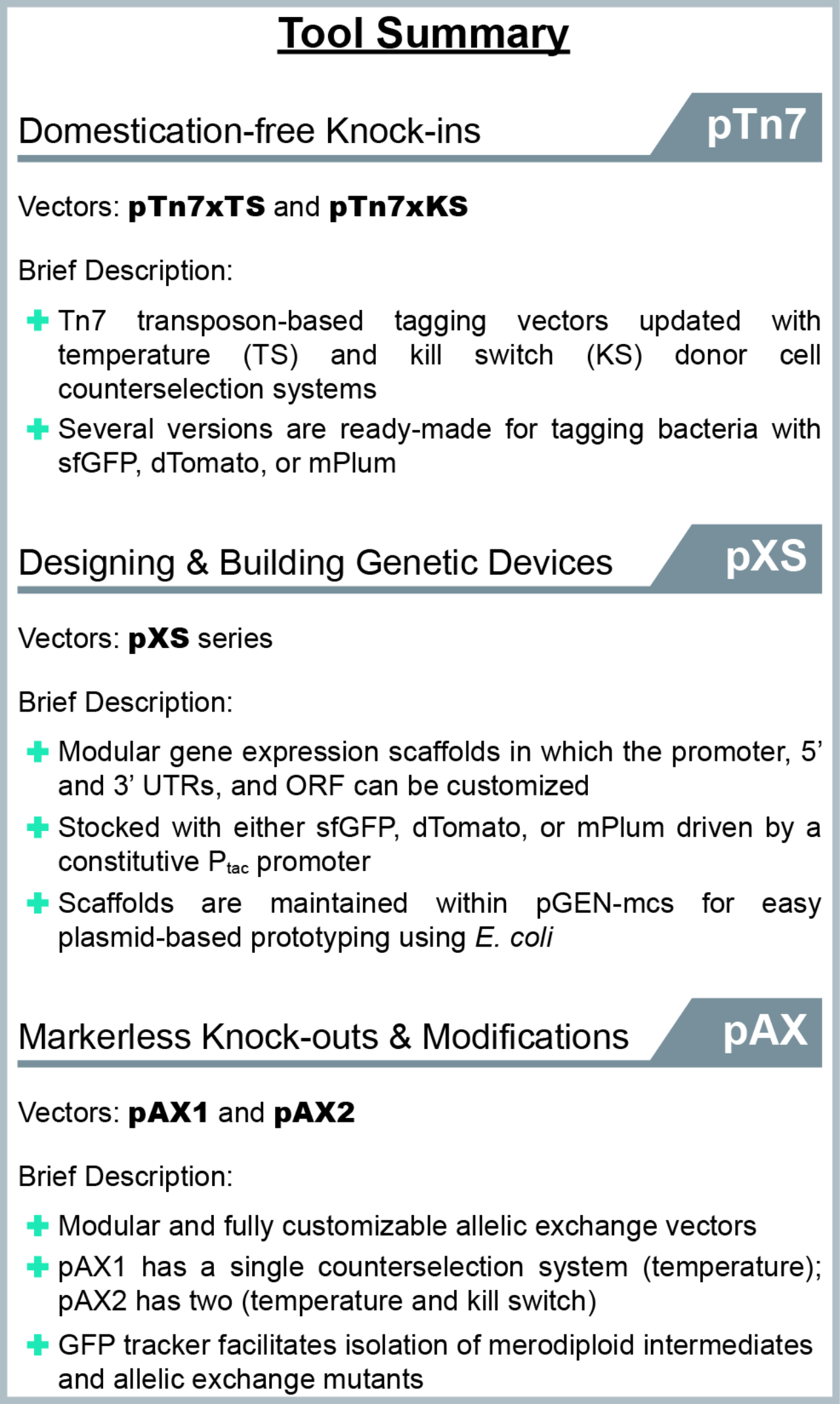
Summary of the genetic tools described in this work. The functions and features of each engineering vector is briefly summarized. Guide is organized based on technique or intended use.

#### Minimizing laboratory-based domestication of wild bacterial isolates

The deleterious nature of domestication is well-documented, yet it is often overlooked and domestication steps are unfortunately routinely performed because of their convenience^23,24^. Domestication is commonly used to improve genetic tractability or to help discern specific bacterial strains from other lineages within complex environments (e.g., the vertebrate intestine, water, or soil). Compensatory mutations can rescue or mask physiological defects associated with domestication in vitro^25^; however, there is no guarantee that critical aspects of symbiont biology, such as those involved in host engagement, are left unperturbed. Therefore, the accurate modeling of symbiotic interactions requires careful attention to preserving natural symbiont behaviors. The incorporation of temperature and kill switch-based domestication-free counterselection systems into a previously described Tn7 tagging vector^27^, which produced pTn7xTS and pTn7xKS (Figure 2), and the novel allelic exchange vectors pAX1 and pAX2 developed in this work (Figure 5), offers ready-made tools for manipulating various symbiotic bacterial lineages in a way that preserves their natural physiology.

#### Achieving broad utility through modularity

The incredible genetic and phenotypic diversity of bacteria challenges cross-lineage compatibility of genetic tools. A major contributing factor to this problem is that many available tools are irreversibly constructed, which impedes the customization of important sequence motifs for different bacteria. We addressed this by building tools with highly modular architectures so that they can be easily reconfigured and are thus molecularly nimble. This feature makes it possible to continuously innovate and build off original designs. For example, expanding on the expression scaffolds within the pXS series of vectors (Figure 3 and Figure 3-Figure Supplement 1), we have engineered more elaborate genetic devices, including reporters of gene expression and genetic switches. Additionally, the modularity of these vectors can be advantageous in situations where rational design is not possible due to the unavailability of suitable sequence elements; libraries in which constructs containing a single variable motif (e.g., a promoter or ribosome binding site) can be readily assembled and screened for optimal activity. While the flexibility of our expression scaffolds is conducive to further engineering, as built they have immediate utility for stably tagging bacteria and thus, facilitating the direct observation of symbionts within their natural host-associated environments. Illustrating this point, the mere fluorescent tagging of the zebrafish symbionts *Vibrio* ZWU0020 and *Aeromonas* ZOR0001 using these tools revealed that each bacterium interacts with the physically dynamic confines of the intestine in distinct ways and that this differential interplay unexpectedly shapes their apparent competition^13^. We also incorporated modularity into the design of the allelic exchange vectors pAX1 and pAX2 (Figure 5 and Figure 5-Figure Supplement 1). Several elements within these vectors can be customized, including the antibiotic resistance genes, the fluorescent merodiploid tracker, and the kill switch. With this modularity, we are exploring alternative utilities of these vectors. For example, swapping the GFP merodiploid tracker for one that encodes a red fluorescent protein has allowed us to engineer GFP fusions at endogenous chromosomal loci. In total, tailorable tools such as those described here offer a way of tuning or customizing functionality to increase the experimental potential of bacterial lineages.

#### Streamlining genetic manipulations with visual screening

To improve the tractability of allelic exchange we used GFP to visually track recombination events (Figure 4). This simple update proved extremely powerful. It allowed merodiploids to be confidently identified and isolated while final mutant derivatives could be screened for with incredible sensitivity, sometimes being found as small subpopulations within merodiploid colonies growing on an agar plate (Figure 4 and Figure 6-Figure Supplements 1-3). The successful manipulation of a previously intractable bacterium (i.e., *Vibrio* ZOR0035) highlights the utility of this approach. Although conventional selection schemes (e.q., based on *sacB*) are adept at recovering mutants that arise at low frequencies due to rarely occurring recombination events, their use is contingent on specific conditions that can be difficult to translate between species or strains. By contrast, our visual screening approach operates freely across lineages and differs from selections in that it allows for the progression of recombination events to be more precisely monitored. As a result, the engineering and isolation of bacterial mutants is more efficient and attainable.

### Demonstrating New Avenues for Research: Comparative Study of Bacterial Biogeography in the Zebrafish Gut

The technical flexibility of the tools and methods developed in this work makes it easier to genetically manipulate many different bacterial lineages in parallel. This functionality greatly facilitates comparative studies aiming to disentangle unique and widely conserved aspects of host-microbe systems. Such investigations are important because many properties of complex host-associated microbial communities, like those comprising human microbiota^54,55^, are extremely variable and remain unexplained. Illustrating the potential for comparative approaches, we exposed several uncharacterized phenomena by probing the colonization patterns and behaviors of multiple bacterial lineages native to the zebrafish intestine.

Upon initial examination, we found discordance between predicted and observed motility phenotypes in vivo. All strains examined carry flagellar genes and tend to be highly motile in vitro, yet several of them (e.g., *Enterobacter* ZOR0014, *Aeromonas* ZOR000l, and *Aeromonas* ZOR0002) display no obvious motility during intestinal colonization (Supplementary Movies 2-4). By contrast, some strains robustly sustain motility in vivo, most notably *Vibrio* ZWU0020, which appears to produce populations almost entirely made up of swimming cells (Supplementary Movie 1). There are several possible explanations for these behaviors that are not necessarily mutually exclusive. For example, they may be the result of bacteria dynamically responding to the intestinal environment and executing strain or lineage-specific colonization strategies. On the other hand, bacteria could be differentially susceptible to some form of host-mediated motility interference, which has recently been documented in mouse models^56,57^. Work focused on distinguishing these interactions is ongoing. Ultimately, this observation highlights the disconnect that can occur among in silico, in vitro, and in vivo approaches for studying bacterial symbioses. Considering how to best capture and interpret mechanistic insights across model systems will be critical to progress.

Our data additionally indicate that some strains display a relatively large amount of variation in growth mode and biogeography between hosts (e.g., *Aeromonas* ZOR000l, *Aeromonas* ZOR0002, and *Enterobacter* ZOR0014), while others do not (e.g., *Plesiomonas* ZOR0011, *Pseudomonas* ZWU0006, and *Vibrio* ZWU0020) (Figure 8). This variance is consistent with earlier reports showing that the structure of bacterial populations within the larval zebrafish gut can be highly dynamic, which is attributable in part to the physical forces of intestinal peristalsis^13,49^. Yet some bacteria—for example, *Vibrio* ZWU0020—remain stable in the face of this perturbation through still undefined mechanisms^13^. Our observations with the *Vibrio* ZWU0020 Δ*pomAB* mutant suggest cell motility is involved.

Most strikingly, we discovered a strong correlation between the dominant growth mode of bacterial populations and their biogeography (Figure 8). While cell behavior is recognized to influence local population structure^58,59^, linking cell aggregation with a global pattern of spatial organization throughout the intestine is unanticipated and profound. A possible explanation for this pattern is that physical properties of the intestinal environment (e.g., its shape and/or peristaltic movement) act to spatially segregate planktonic and aggregated cells. Alternatively, bacteria may toggle between different growth modes in response to spatial cues generated by physiologically distinct regions along the length of the intestine. Going forward, a major objective will be to dissect the potential mechanisms of this relationship and importantly, understand how it generalizes both within and across host–microbe systems.

### Outlook

Elucidating the rules that govern the assembly and function of bacterial symbioses requires studying a wide range of bacterial symbiont lineages^1,2^. Whether abundant, rare, divergent, or closely related, all potentially hold clues to how host-microbe systems work and how they can be exploited for biotechnology applications—from boosting food production to treating human disease. As more symbiotic relationships are uncovered and new model systems come online, the continued design and modernization of genetic approaches for streamlined manipulation of diverse bacterial lineages will be paramount. Although symbiotic bacteria were the primary subject for tool development in this work, the approaches we have described are equally applicable to the study of free-living environmental bacteria. Importantly, the growing appreciation for the ubiquity of bacterial symbioses and their far-reaching influence on the lives of plants and animals is inspiring a highly cross-disciplinary generation of microbiologists with mixed and varied backgrounds. Therefore, it will be beneficial to work toward standardized tools and methods that foster the rigorous and accurate investigation of symbiont biology.

## MATERIALS AND METHODS

### Animal care

All experiments with zebrafish were done in accordance with protocols approved by the University of Oregon Institutional Animal Care and Use Committee and following standard protocols^60^.

### Gnotobiology

Wild-type (AB × TU strain) zebrafish were derived germ-free (GF) and colonized with bacterial strains as previously described with slight modification^61^. Briefly, fertilized eggs from adult mating pairs were harvested and incubated in sterile embryo media (EM) containing ampicillin (100μg/ml), gentamicin (10μg/ml), amphotericin B (250ng/ml), tetracycline (1μg/ml), and chloramphenicol (1μg/ml) for ~6h. Embryos were then washed in EM containing 0.1% polyvinylpyrrolidone–iodine followed by EM containing 0.003% sodium hypochlorite. Sterilized embryos were distributed into T25 tissue culture flasks containing 15ml sterile EM at a density of one embryo per ml and incubated at 28-30°C prior to bacterial colonization. Embryos were sustained on yolk-derived nutrients and not fed during experiments. For bacterial mono-association, bacterial strains were grown overnight in LB liquid media with shaking at 30°C, and prepared for inoculation by pelleting 1ml of culture for 2 min at 7,000 × g, and washing once in sterile EM. Bacterial strains were individually added to the water column of single flasks containing 4-day old larval zebrafish at a final density of ~10^6^ bacteria/ml. Bacterial colonization patterns were assessed 24h later by live imaging of three separate 5-day old zebrafish hosts per bacterial strain. Three animals were determined to be adequate for capturing general colonization features of each bacterial strain based on at least two previous independent qualitative assessments of colonization patterns.

### Bacterial strains and culture

All wild and recombinant bacterial strains used or created in this study are listed in Supplementary File 1. Archived stocks of bacteria are maintained in 25% glycerol at −80°C. Prior to manipulations or experiments, bacteria were directly inoculated into 5ml Luria-Bertani (LB) media (10g/L NaCl, 5g/L yeast extract, 12g/L tryptone, 1g/L glucose) and grown for ~16h (overnight) shaking at 30°C, except for *E. coli* HS, which was grown at 37°C. For growth on solid media, tryptic soy agar was used. 10μg/ml gentamicin was used to select recombinant strains tagged with the Tn7 transposon, which was modified to carry a gentamicin resistance gene. When selecting merodiploid intermediates made using pAX1 or pAX2, which carry resistance to both gentamicin and chloramphenicol, either 10μg/ml gentamicin or 5μg/ml chloramphenicol was used. Selection of rifampicin-domesticated variants was done using 100μg/ml rifampicin.

### Molecular techniques and reagents

*E. coli* strains used for molecular cloning and conjugation, and plasmids used or created during this work are listed in Supplementary File 2. *E. coli* were typically grown in 5ml LB liquid media at 30°C or 37°C with shaking in the presence of appropriate antibiotic selection to maintain plasmids. For propagation on solid media, LB agar was used. Antibiotic concentrations used were as follows: 100μg/ml ampicillin, 20μg/ml chloramphenicol, 10μg/ml gentamicin, and 10μg/ml tetracycline. Supplementary File 3 lists all DNA primers used for polymerase chain reactions (PCR), which are organized based on their “Wiles Primer” (WP) number. Unless specifically noted, standard molecular techniques were applied and reagents were used according to manufacturer instructions. Restriction enzymes and other molecular biology reagents for PCR and nucleic acid modifications were obtained from New England BioLabs (Ipswich, MA). Various kits for plasmid and PCR amplicon purification were obtained from Zymo Research (Irvine, CA). The Wizard Genomic DNA Purification Kit (Promega, Madison, WI) was used for isolating bacterial genomic DNA. DNA oligonucleotides for PCR were synthesized by Integrated DNA Technologies (Coralville, IA). Sanger sequencing was done by Sequetech (Mountain View, CA). Custom gene synthesis was done by GenScript (Piscataway, NJ). A Leica MZ10 F fluorescence stereomicroscope with 1.0x, 1.6x, and 2.0x objectives and Leica DFC365 FX camera were used for screening and imaging fluorescent bacterial colonies (Leica, Wetzlar, Germany). Images were captured and processed using standard Leica Application Suite software and ImageJ^62^. Nucleotide sequences of 16S rRNA genes used for phylogenetic analysis are provided in Supplementary File 4, and were obtained via “The Integrated Microbial Genomes & Microbiome Samples” (IMG/M) website (https://img.jgi.doe.gov/m/)^63^ or the RNAmmer web tool^64^. 16S rRNA sequences were aligned using Clustal Omega^65^ and an unrooted phylogenetic tree was drawn using the Phylodendron web tool (http://iubio.bio.indiana.edu:7131/treeapp/treeprint-form.html).

### Plasmid construction

The plasmid-based tools developed in this work have been deposited at Addgene (Cambridge, MA), along with their sequences (https://www.addgene.org/). Supplementary File 5 contains annotated nucleotide sequences of select genetic parts that were used to build plasmids and gene expression scaffolds. Details on how plasmids were specifically constructed are provided in Supplementary File 6.

### Domestication-free Tn7 tagging using pTn7xTS and pTn7xKS

A detailed Tn7 tagging protocol based on pTn7xTS and pTn7xKS—which includes optimization and troubleshooting steps, and notes on strain-specific procedures—is provided in Supplementary File 7. Generally, and as outlined in Figure 2, triparental conjugation was performed between a single target bacterial strain, an *E. coli* SM10 donor strain carrying the transposase-containing pTNS2 helper plasmid, and an *E. coli* SM10 donor strain carrying either a pTn7xTS or pTn7xKS domestication-free tagging vector. Prior to mating, bacteria were prepared by subculturing them to an approximate optical density of 0.4-0.6 at 600nm in LB media with required antibiotics and at the appropriate growth temperature. Cells were then combined 1:1:1 (500μl each), washed once by centrifugation and aspiration in 1ml LB media or 0. 7% NaCl, and suspended in a final 25μl volume of the same media used for washing. Next, the concentrated mating mixture was transferred to a 25mm-wide 0.45μm filter disc (EMD Millipore, Billerica MA; product #HAWP02500) that had been placed on top of a TSA plate. Once the mating mixture dried, the plate was incubated at 30°C for 3-5h. After incubation, the filter disc was placed in 1ml 0.7% NaCl within a 50ml conical tube and bacteria were dislodged by vortexing and pipetting. In cases where a pTn7xTS-based vector was used, 100μl of the bacterial suspension was spread onto a TSA plate containing gentamicin and incubated overnight at 37°C to select for recombinants. To ensure the recovery of low frequency recombinants, the remaining 900μl of the suspension was pelleted by centrifugation, suspended in 100μl 0.7% NaCl, and plated in the same way. In cases where a pTn7xKS-based vector was used, 100μl of the bacterial suspension was spread onto a TSA plate containing gentamicin and 1mM isopropyl-β-D-thiogalactoside (IPTG), and incubated overnight at 30°C. The remaining 900μl was prepared as above, plated on TSA with gentamicin and IPTG, and incubated at 30°C.

The following day, putative recombinant target bacteria were colony-purified by streaking on TSA without antibiotic selection at 30°C. Of note, when recombinant bacteria are tagged with a gene encoding a fluorescent protein, performing colony-purification in the absence of antibiotic selection followed by visual screening of fluorescence is a convenient way to verify that the Tn7 transposon has chromosomally integrated and the tagging vector has been lost. Purified clones were picked, cultured in LB media containing gentamicin, and genotyped by PCR to verify correct insertion of the Tn7 transposon at the *attTn7* site. The universal primer WP11, which anneals within the Tn7R region of the Tn7 transposon, was used with a species-specific primer that anneals to an adjacent chromosomal sequence within the 3’ end of the *glmS* gene to generate a small (~250bp) amplicon if the transposon is present. Species-specific primers used were as follows: *Vibrio* Z0R0018, WP50; *Vibrio* Z0R0035, WP51; *Vibrio* Z0R0036, WP12; *Vibrio* ZWU0020, WP12; *Aeromonas* Z0R0001, WP49; *Aeromonas* Z0R0002, WP52; *Pseudomonas* ZWU0006, WP256; *Acinetobacter* Z0R0008, WP259; *Enterobacter* Z0R0014, WP257; *Plesiomonas* Z0R0011, WP260; *E. coli* HS, WP150; *Aeromonas* HM21, WP49; *Shewanella* MR-1, WP48.

### Generation of markerless deletions via allelic exchange using pAX1 and pAX2

A detailed protocol for carrying out allelic exchange using pAX1 and pAX2—which includes optimization and troubleshooting steps, and notes on strain-specific procedures—is provided in Supplementary File 8. Briefly, and as summarized in Figure 4, allelic exchange cassettes for mediating markerless deletion of target genetic loci (i.e., the *pomAB* locus of *Vibrio* ZWU0020 and *Aeromonas* ZOR0001) were generated through splice by overlap extension and inserted into a pAX-based vector. Next, diparental conjugation was performed between a single target bacterial strain (i.e., *Vibrio* ZWU0020 or *Aeromonas* ZOR0001) and an *E. coli* SM10 donor strain carrying the assembled allelic exchange vector. Prior to mating, bacteria were prepared by subculturing them to an approximate optical density of 0.4-0.6 at 600nm in LB media with required antibiotics and at the appropriate growth temperature. Cells were then combined 1:1 (750μl each), washed once by centrifugation and aspiration in 1ml LB media or 0.7% NaCl, and suspended in a final 25μl volume of the same media used for washing. Next, the concentrated mating mixture was transferred to a 25mm-wide 0.45μm filter disc that had been placed on top of a TSA plate. Once the mating mixture dried, the plate was incubated at 30°C for 3-5h. After incubation, the filter disc was placed in 1ml 0.7% NaCl within a 50ml conical tube and bacteria were dislodged by vortexing and pipetting. For the generation of *Vibrio* ZWU0020 Δ*pomAB*, which employed a pAX1-related vector, 100μl of the bacterial suspension was spread onto a TSA plate containing gentamicin and incubated overnight at 37°C to select for merodiploids. The remaining 900μl of the suspension was pelleted by centrifugation, suspended in 100μl 0.7% NaCl, and plated in the same way to ensure recovery of rare recombinants. For the generation of *Aeromonas* ZOR0001 Δ*pomAB*, which employed a pAX2-based vector, 100μl of the bacterial suspension was spread onto a TSA plate containing gentamicin and 10ng/ml anhydrotetracycline (aTc), and incubated overnight at 30°C. The remaining 900μl was prepared as above, plated on TSA with gentamicin and aTc, and incubated at 30°C.

The following day, colonies of putative merodiploid target bacteria that were expressing the GFP tracker were purified by streaking on TSA without antibiotic selection at 30°C. This purification step also served to verify that the allelic exchange vector had integrated into the chromosome. Purified clones were picked, cultured in LB media containing gentamicin to maintain their merodiploid state, and archived as a frozen stock. To screen for second recombination events, merodiploids were cultured overnight in LB media without antibiotic selection and spread onto several TSA plates, again without antibiotic selection, at a density that allowed ~100-200 discrete colonies to form. Colonies exhibiting partial or complete loss of GFP expression were purified by streaking on TSA at 30°C. Putative mutants were screened and genotyped by PCR using primers that flanked the modified locus, which produced two differently sized amplicons that represented the wild-type and mutant alleles. Primers WP163 and WP164 were used to genotype *Vibrio* ZWU0020 Δ*pomAB* mutants and primers WP192 and WP195 were used to genotype *Aeromonas* Z0R0001 Δ*pomAB* mutants.

### In vitro growth measurements

In vitro growth of bacterial strains was assessed using the FLUOstar Omega microplate reader (BMG LABTECH, Offenburg, Germany). Prior to growth measurements, bacteria were grown overnight in 5ml LB media at 30°C with shaking. Cultures were diluted 1:100 into fresh LB media and dispensed in quadruplicate (i.e., four technical replicates) (200μl/ well) into a sterile 96 well clear flat bottom tissue culture-treated microplate (Corning, Corning, NY; product #3585). Absorbance measurements at 600nm were then recorded every 30 minutes for 16 hours (or until stationary phase) at 30°C with shaking. Growth measurements were repeated at least two independent times for each strain (i.e., two biological replicates) with consistent results. Data were exported and graphed using GraphPad Prism 6 software.

### Swim motility assays

Prior to the assessment of swimming motility, bacteria were grown overnight in 5ml LB media at 30°C with shaking. 1ml of bacterial culture was then washed by centrifuging cells at 7,000xg for 2 minutes, aspirating media, and suspending in 1ml 0.7% NaCl. This centrifugation/ aspiration wash step was repeated once more and bacteria were suspended in a final volume of 1ml 0.7% NaCl. 1μl of washed bacterial culture was then inoculated into a TSA plate containing 0.2% agar (30g/L tryptic soy broth and 2g/L bacto agar). Swim plates were incubated at 30°C for 5-7h and imaged on a Gel Doc XR+ Imaging System (Bio-Rad, Hercules, CA). Motility assays were repeated at least two independent times (i.e., two biological replicates) with consistent results.

### Spot tests

*E. coli* SM10 donor cells carrying vectors that contain temperature and/or kill switch-based post-conjugation counterselection systems were grown overnight in LB media with required antibiotics and at the appropriate growth temperatures. For assessing temperature-based counterselection, ten-fold serial dilutions were made on TSA plates containing gentamicin and incubated overnight at 30°C or 37°C. For assessing kill switch-based counterselection, ten-fold serial dilutions were made on TSA plates containing gentamicin +/− 1mM IPTG (in the case of pTn7xKS) or 10ng/ml aTc (in the case of pAX2) and incubated overnight at 30°C. Plates were imaged on a Bio-Rad Gel Doc XR+ Imaging System. All spot tests were performed at least two independent times (i.e., two biological replicates), each including at least two technical replicates, with consistent results.

### Live Imaging

Live larval zebrafish were imaged using a home-built light sheet fluorescence microscope described in detail elsewhere^49,66^. In brief, a thin sheet of laser light is obtained by rapidly scanning the excitation beam with a galvanometer mirror. Fluorescence emission is captured by an objective lens mounted perpendicular to the sheet. 3D images are obtained by translating the sample along the detection axis. The entire volume of the intestine (approximately 1200x300x150 microns) is imaged in four sub-regions that are computationally registered after acquisition. Total acquisition time of a single intestine is less than 1 min with 1-micron steps between planes. For all images, the exposure time was 30ms and the excitation laser power was 5mW prior to entering the imaging chamber.

### Image Analysis

Bacterial abundances and locations were estimated using the analysis pipeline described in^49^. In brief, we identify individual cells in 3D using a wavelet filtering-based algorithm^67^ and identify multicellular aggregates using a graph-cut segmentation algorithm^68^. The number of cells in an aggregate is estimated by dividing the total aggregate intensity by the average intensity of individual bacteria. Individuals detected within an aggregate are discarded. One-dimensional population distributions are obtained by dividing the intestine into 5-micron bins constructed down the length of the intestine along a manually drawn line and assigning the centroid of each detected object to a bin. Global population centers are computed as the center of mass of this 1D distribution. Of note, this analysis pipeline was originally developed and optimized for a different strain not imaged here^49^ and its performance on the 8 present strains has not been rigorously assessed. Based on manual inspection and analysis, we estimate an uncertainty of at most 10% for the planktonic fraction and for the population center of mass, which is certainly more than adequate to detect the global trends we report.

## ACKNOWLEDGEMENTS

We would like to thank numerous members of the Guillemin lab for their willingness to test and provide feedback on the genetic tools developed in this work, particularly Dr. Annah Rolig and Dr. Cathy Robinson. We would also like to thank Dr. Andrew Camilli for his generosity in sharing overlap extension and allelic exchange protocols. This project was supported through the M.J. Murdock Charitable Trust and an award from the Kavli Microbiome Ideas Challenge, a project led by the American Society for Microbiology in partnership with the American Chemical Society and the American Physical Society and supported by The Kavli Foundation. Work was also supported by the National Science Foundation under Awards 0922951 and 1507115. Authors received funding from the National Institutes of Health (NIH, http://www.nih.gov/), P50GM09891 to KG, F32AI112094 to TJW, and T32GM007759 to BHS. The funders had no role in study design, data collection and analysis, decision to publish, or preparation of the manuscript.

## FIGURE SUPPLEMENT AND SUPPLEMENTARY MOVIE LEGENDS

**Figure 2-Figure Supplement 1.**
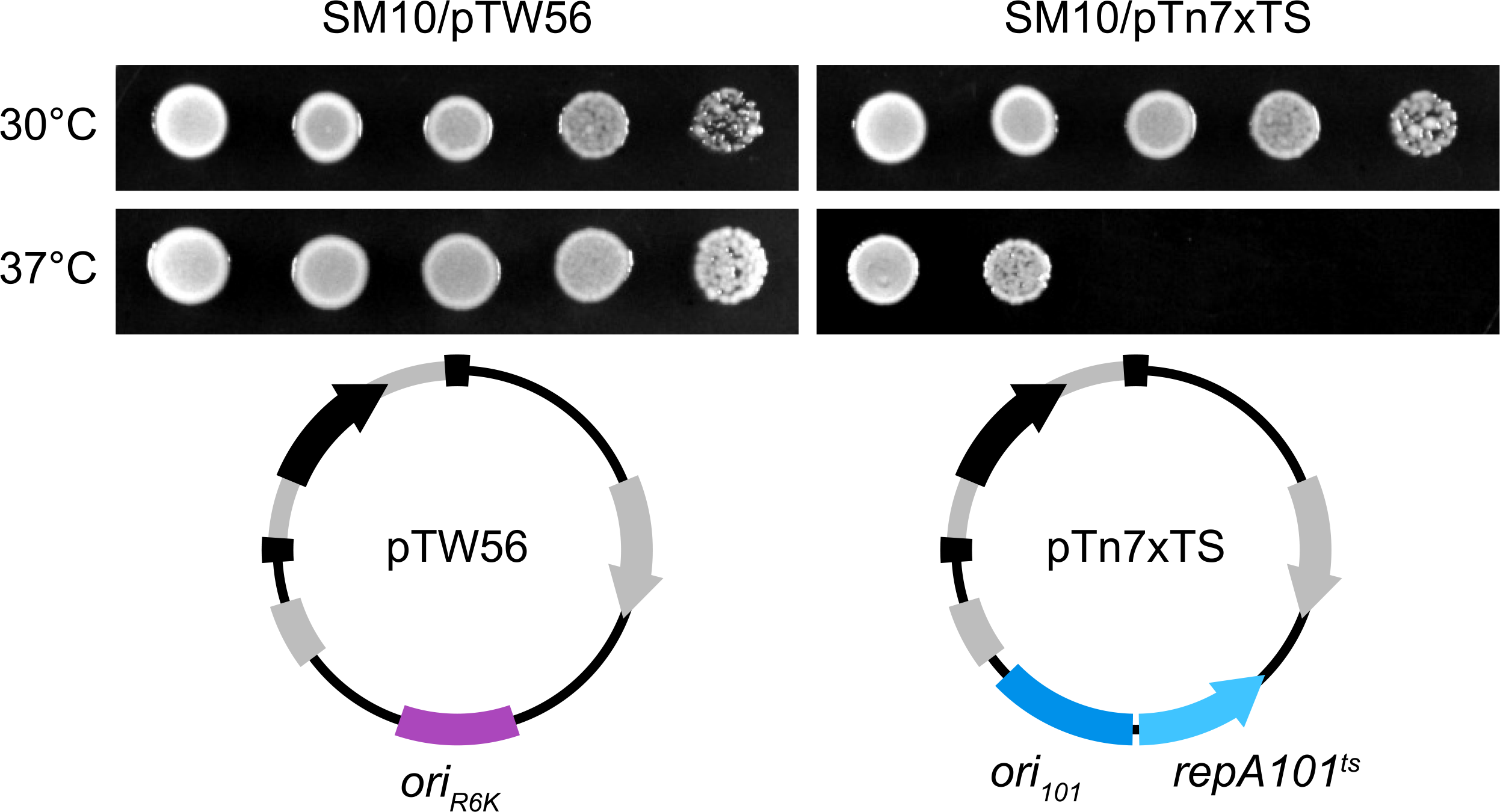
Spot tests demonstrating temperature-based counterselection of *E. coli* SM10 donor cells using the Tn7-taging vector pTn7xTS. Ten-fold serial dilutions of *E. coli* SM10 carrying either the control vector pUC18R6KT-mini-Tn7T-GM (pTW56) or its temperature-sensitive derivative pTn7xTS were plated on tryptic soy agar containing 10μg/ml gentamicin and cultivated overnight at 30°C or 37°C. Functional differences between each vector are highlighted in vector maps. *ori*_*R6K*_, *pir*-dependent origin of replication; *ori*_*101*_/*repA*101^ts^, temperature-sensitive origin of replication.

**Figure 2-Figure Supplement 2.**
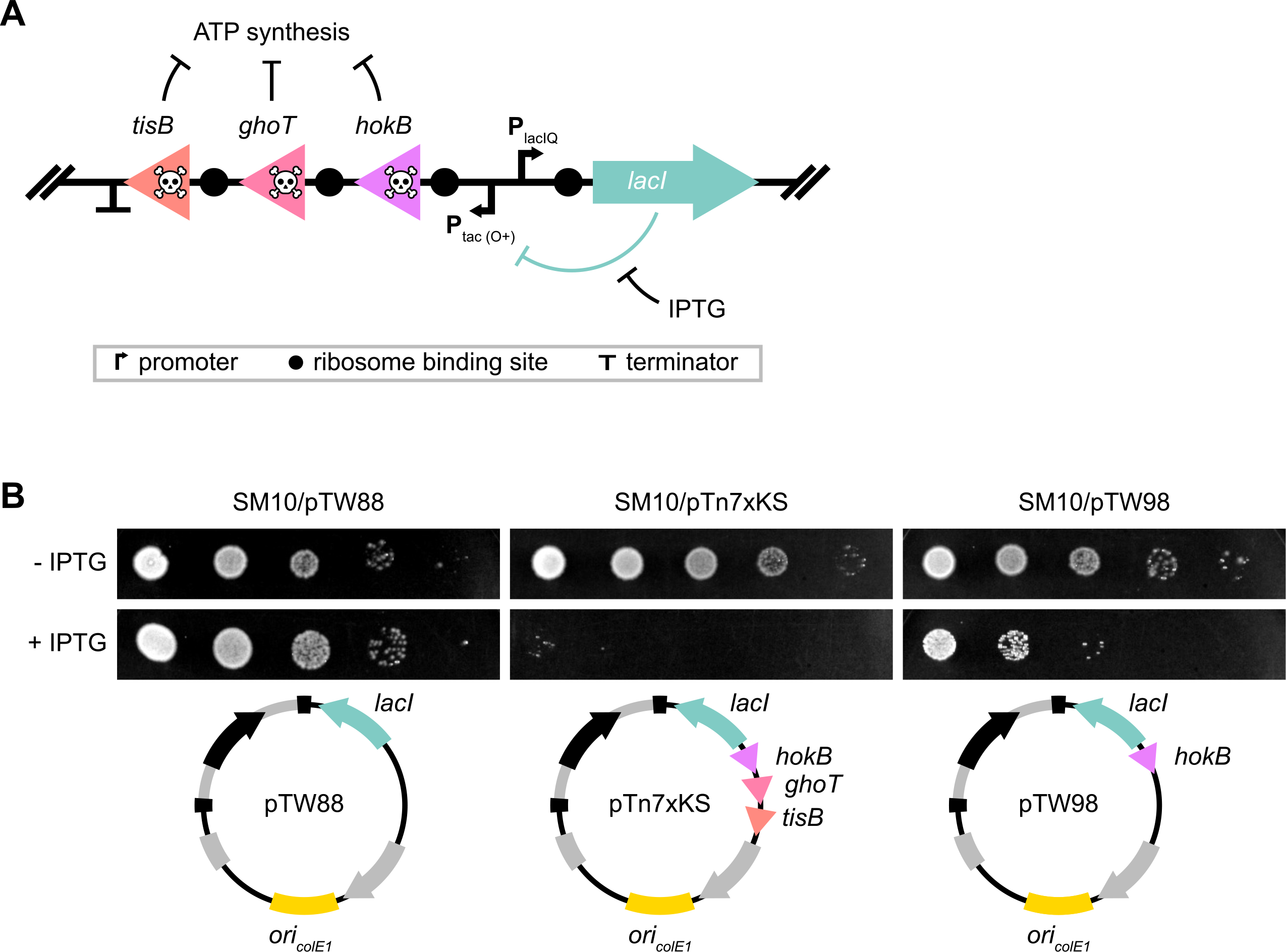
Spot tests demonstrating kill switch-based counterselection of *E. coli* SM10 donor cells using the Tn7-taging vector pTn7xKS. (**A**) Kill switch design. Expression of the three toxin-encoding genes *hokB, ghoT*, and *tisB*—which impair ATP synthesis—is controlled by the LacI-repressible promoter P_tac_ (O+). “O+” indicates presence of the lac operator sequence in the promoter. Tight regulation of toxin genes is achieved by using the P_laclQ_ promoter variant to control transcription of *lacI*, which drives ten-fold higher expression over native P_lacl_. The allolactose analogue isopropyl-β-D-thiogalactoside (IPTG) is used for kill switch induction. (**B**) Ten-fold serial dilutions of *E. coli* SM10, carrying pUC18T-mini-Tn7T-GM (pTW54) derivatives that either contain a kill switch with no toxin genes (pTW88), three toxin genes (pTn7xKS), or a single toxin gene (pTW98), were plated on tryptic soy agar containing 10μg/ml gentamicin +/− 1mM IPTG and cultivated overnight at 30°C. Functional differences between each vector are highlighted in vector maps. *ori_ColE1_*, high copy number origin of replication.

**Figure 3-Figure Supplement 1.**
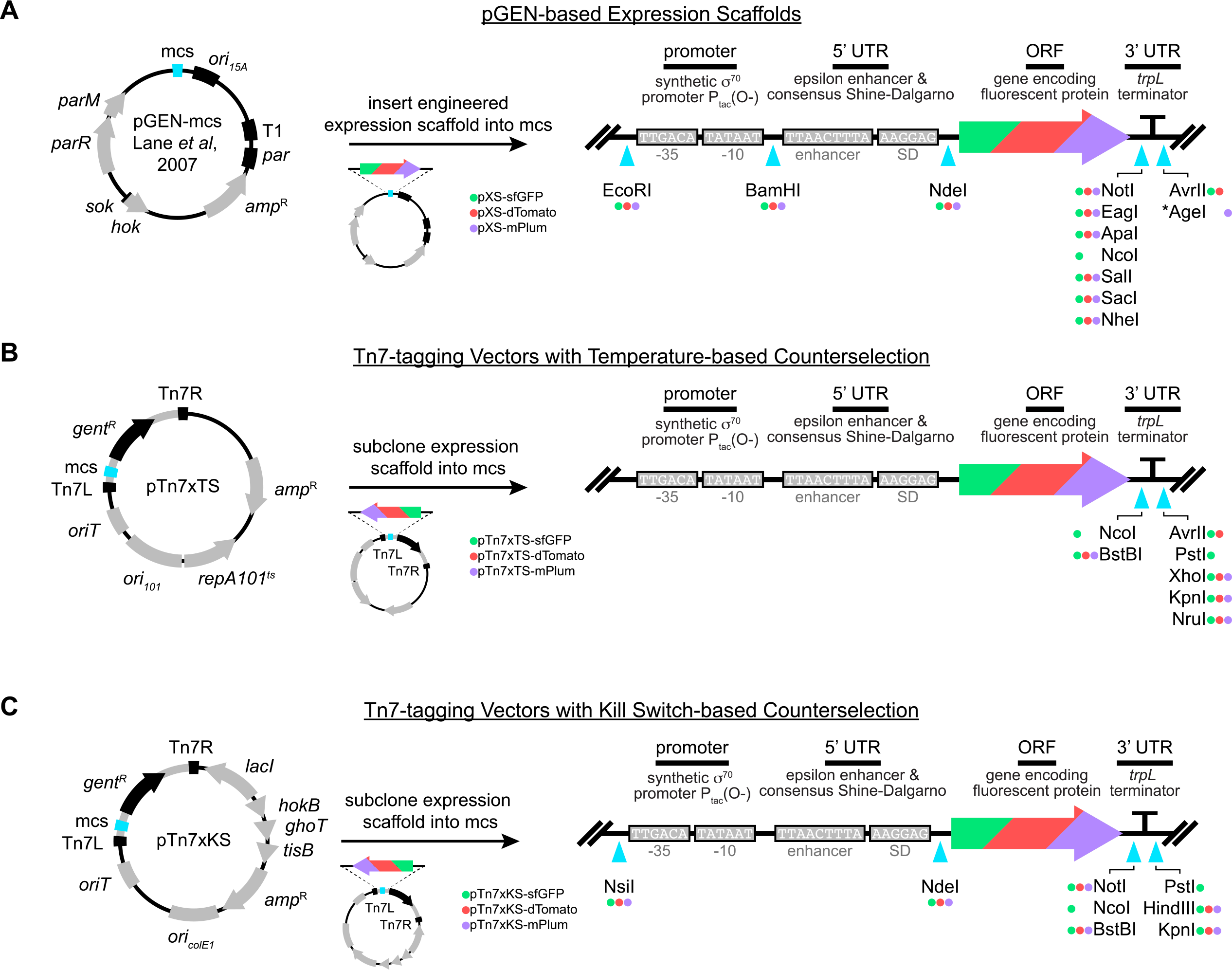
Design features of expression scaffolds and modernized Tn7-tagging vectors. (**A**) The previously described high-retention pGEN-mcs vector was chosen for maintaining, modifying, and prototyping expression scaffolds. Scaffolds encoding either sfGFP (pXS-sfGFP, green circle), dTomato (pXS-dTomato, red circle), or mPlum (pXS-mPlum, purple circle) were built into the multiple cloning site (mcs). Scaffold orientations relative to vector are noted. As in Figure 3, a scaffold diagram highlights the interchangeable sequence motifs and cyan arrowheads below mark the locations of restriction sites for making modifications. Presence of green, red, or purple circles next to restriction enzyme names indicate the availability of that site within each respective vector. *It should be noted that the AgeI enzyme cuts 190bp into the 15A origin of replication. Other pGEN-mcs features: *ori*_*15A*_, broad host range origin of replication; T1, transcriptional terminator; *par*, partitioning system from pSC101; *amp*^R^, ampicillin resistance gene; *hok/sok*, toxin/antitoxin system; *parRM*, centromere-like partitioning system. (**B**) Expression scaffolds were subcloned into the mcs of the temperature-sensitive Tn7-tagging vector pTn7xTS (described in Figure 2A). The relative orientation of the scaffolds is noted. Restriction site availability for each vector encoding either GFP (pTn7xTS-sfGFP, green circle), dTomato (pTn7xTS-dTomato, red circle), or mPlum (pTn7xTS-mPlum, purple circle) is outlined as in **A**. (**C**) Expression scaffolds were subcloned into the mcs of the kill switch containing Tn7-tagging vector pTn7xKS (described in Figure 2C). The relative orientation of the scaffolds is noted. Restriction site availability for each vector encoding either GFP (pTn7xKS-sfGFP, green circle), dTomato (pTn7xKS-dTomato, red circle), or mPlum (pTn7xKS-mPlum, purple circle) is outlined as in **A**.

**Figure 3-Figure Supplement 2.**
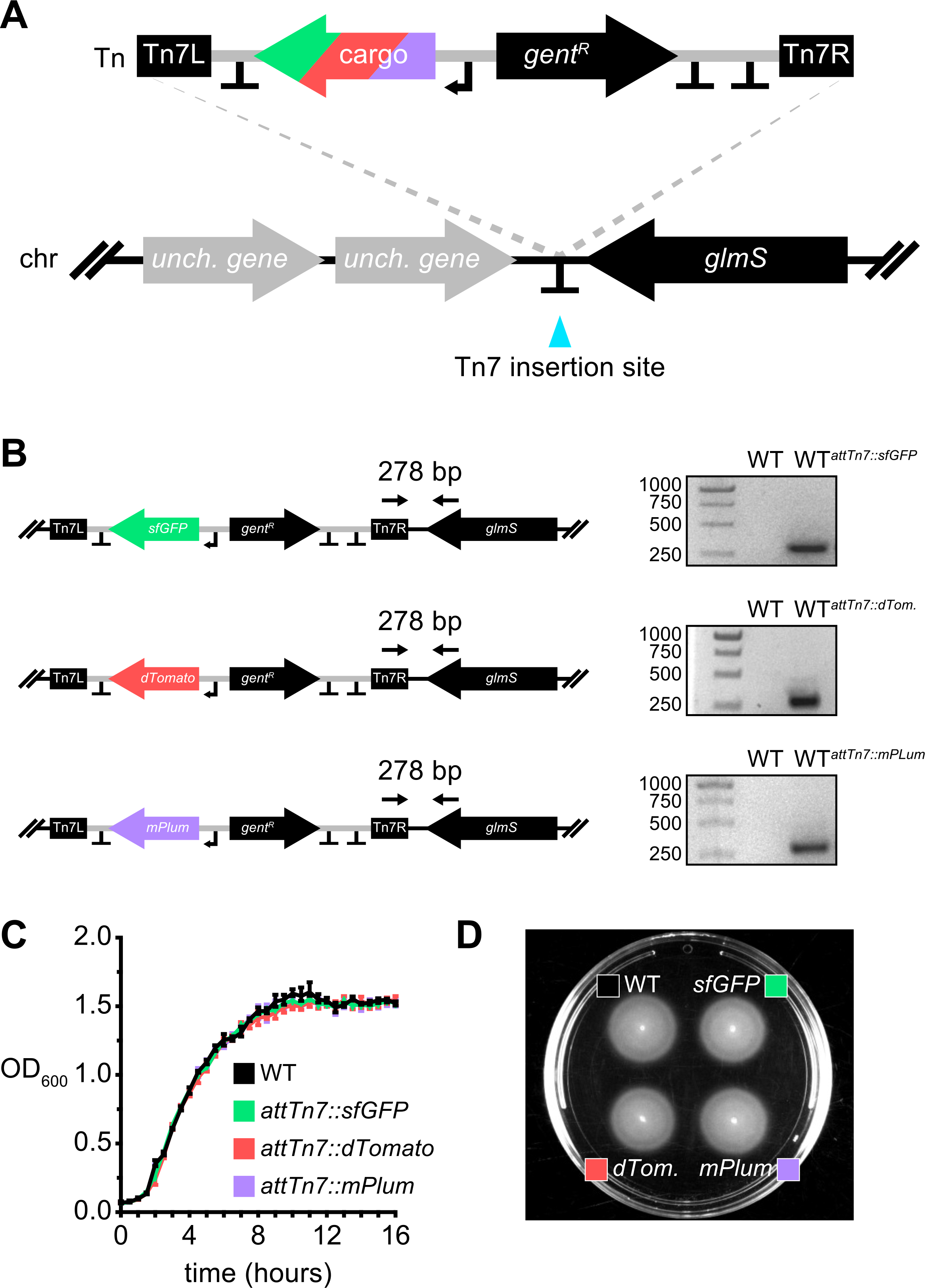
Domestication-free tagging of *Vibrio* ZWU0020. (**A**) Diagram shows the conserved *attTn7* insertion site for Tn7 (Tn) downstream of the *glmS* gene within *Vibrio* ZWU0020’s chromosome (chr). Cargo comprises one of three genes encoding either sfGFP, dTomato, or mPlum. “T” denotes transcriptional terminators; unch., uncharacterized. (**B**) PCR-confirmation of Tn7 insertion variants carrying different fluorescent markers made using temperature and kill switch-based tagging vectors (i.e., pTn7xTS and pTn7xKS, respectively). Black arrows above gene diagrams mark primer annealing sites in addition to the expected amplicon size for successful insertion events. Representative DNA gels showing genotyping results are provided to the right of each diagram. Wild-type ZWU0020 (WT) produces no amplicon because the transposon is not present. (**C**) Plotted is the average optical density at 600nm (OD_600_) vs. time (hours) of WT ZWU0020, and its undomesticated fluorescently tagged derivatives (*attTn7*∷*sfGFP*, *attTn7*∷*dTomato*, and *attTn7*∷*mPlum*), during shaking growth in LB broth at 30°C. Range bars are based on four technical replicates. (**D**) Swim motility WT ZWU0020 and the indicated fluorescently tagged variants from **C** in 0.2% tryptic soy agar at 30°C.

**Figure 5-Figure Supplement 1.**
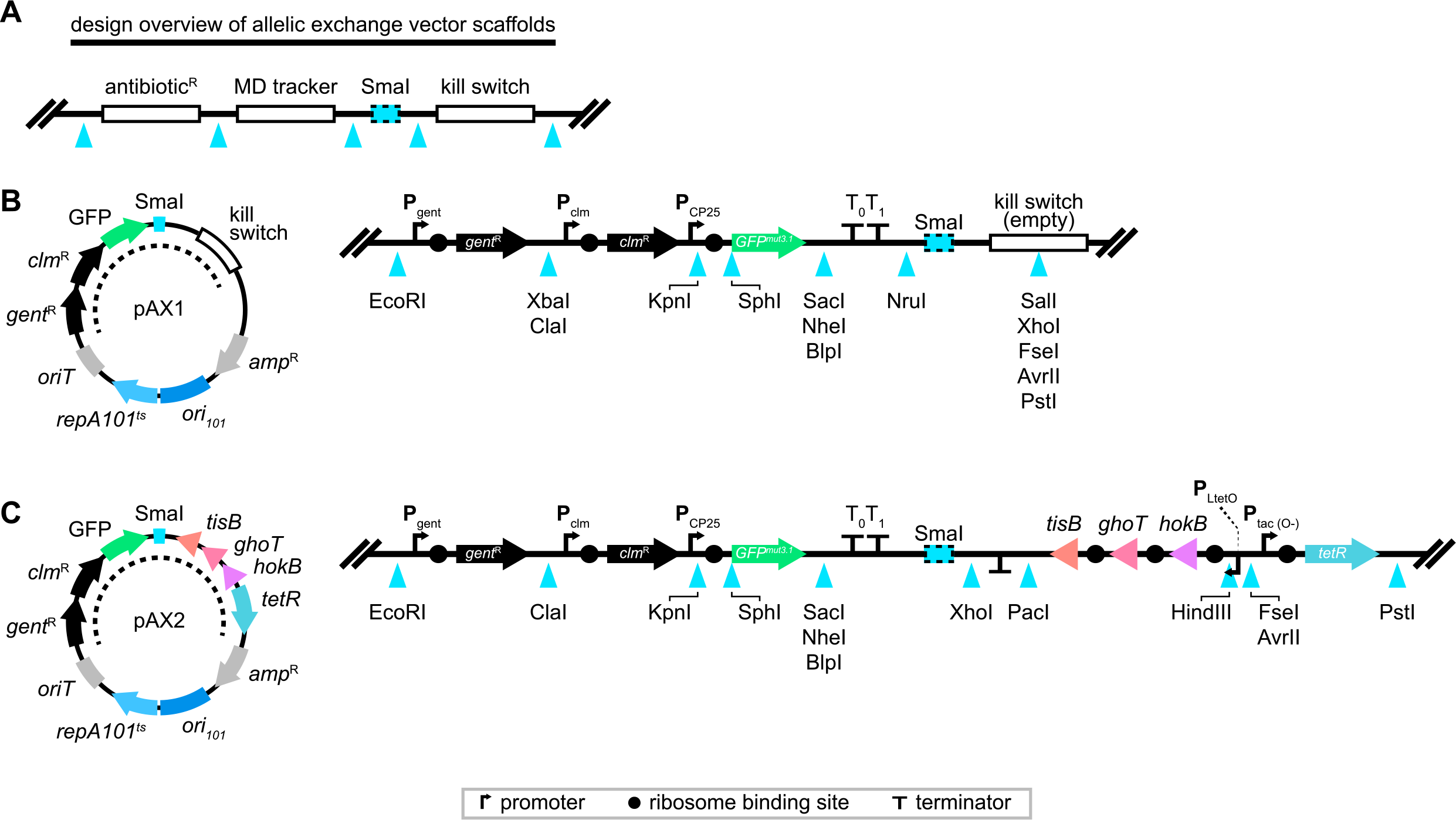
Design features of customizable allelic exchange vectors. (**A**) Molecular scaffolds that hold antibiotic resistance genes, a merodiploid (MD) tracker, and a domestication-free kill switch system are partitioned by restriction sites (cyan arrowheads). A SmaI restriction site, which produces blunt ends when cleaved, serves as a flexible point of entry for cloned allelic exchange cassettes. Full vector assemblies are shown for (**B**) pAX1 and (**C**) pAX2. The hashed line within each vector schematic highlights the scaffold regions shown to the right, which list the locations of available restriction sites for engineering and provide element details. Terminators T_0_ and T_1_ prevent transcriptional read-through of the SmaI cloning site and interference of kill switch elements. *ori*_*101*_/*repA*101^ts^, temperature-sensitive origin of replication; *oriT*, origin of transfer; *gent*^R^, gentamicin resistance gene; *clm*^R^, chloramphenicol resistance gene; *amp*^R^, ampicillin resistance gene; P_gent_, *gent* promoter; P_clm_, *clm* promoter; P_CP25_, synthetic constitutive promoter; GFP^mut3.1^, GFP variant; *tisB, ghoT*, and *hokB* encode toxic peptides; *tetR* encodes the repressor TetR; P_LtetO_, synthetic TetR-repressible promoter; P_tac_ (0-), synthetic constitutive promoter without lac operator sequence.

**Figure 5-Figure Supplement 2.**
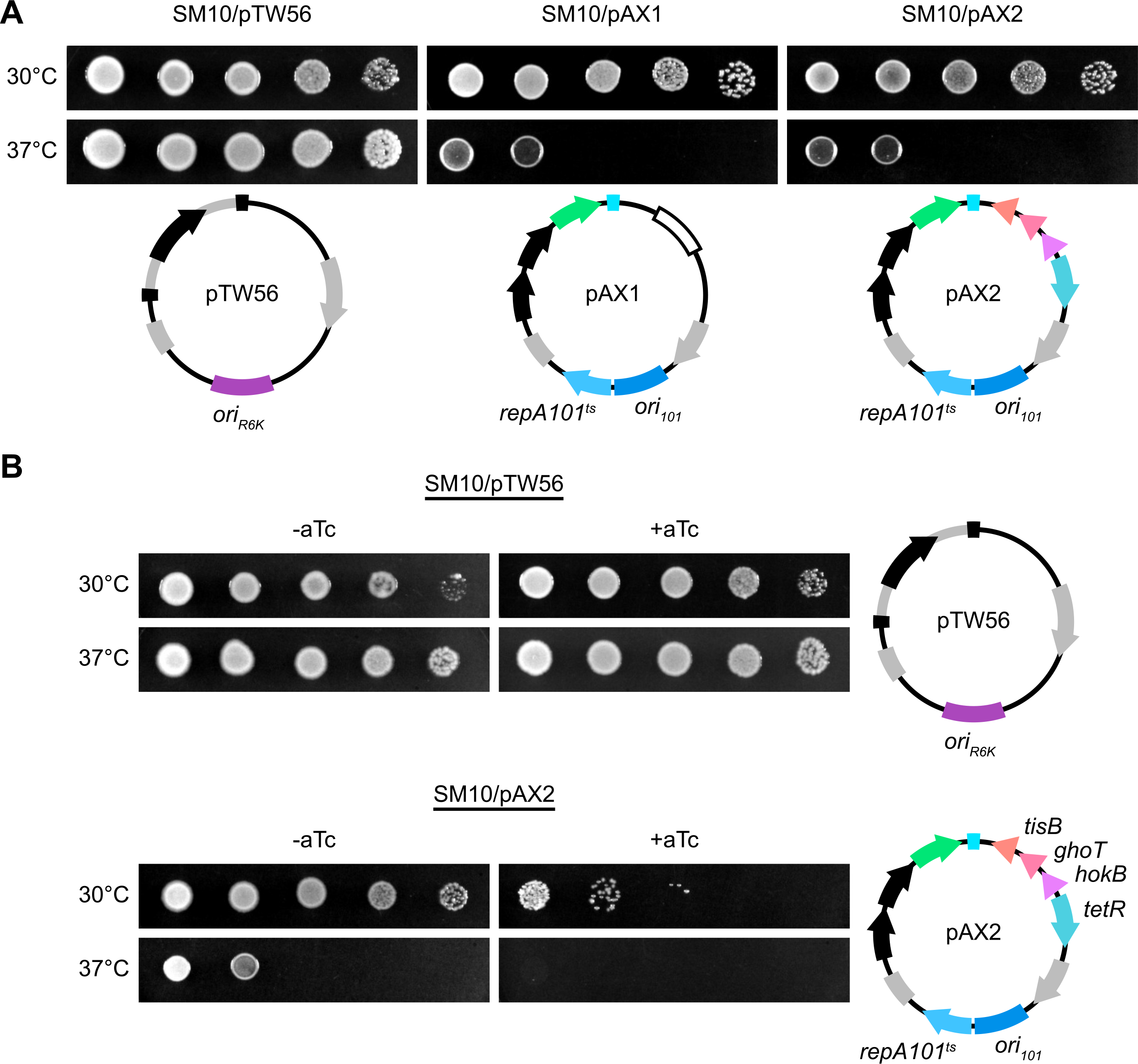
Spot tests demonstrating temperature and kill switch-based counterselection of *E. coli* SM10 donor cells using the allelic exchange vectors pAX1 and pAX2. Ten-fold serial dilutions of *E. coli* SM10 carrying either pAX1 or pAX2 were plated on tryptic soy agar (TSA) containing 10μg/ml gentamicin and cultivated overnight at 30°C or 37°C. Results obtained with SM10/pTW56, which are also presented in Figure 2-Figure Supplement 1, are provided as a reference control. Functional differences between each vector are highlighted. *ori*_*R6K*_, *pir*-dependent origin of replication; *ori*_*101*_/*repA*101^ts^, temperature-sensitive origin of replication. (**B**) Ten-fold serial dilutions of *E. coli* SM10 carrying either pTW56 or pAX2 were plated on TSA containing 10μg/ml gentamicin +/− 10ng/ml anhydrotetracycline (aTc) and cultivated overnight at 30°C or 37°C. As in **A**, functional differences between each vector are highlighted in vector maps. *tisB, ghoT*, and *hokB* encode toxic peptides; *tetR* encodes the repressor TetR.

**Figure 6-Figure Supplement 1.**
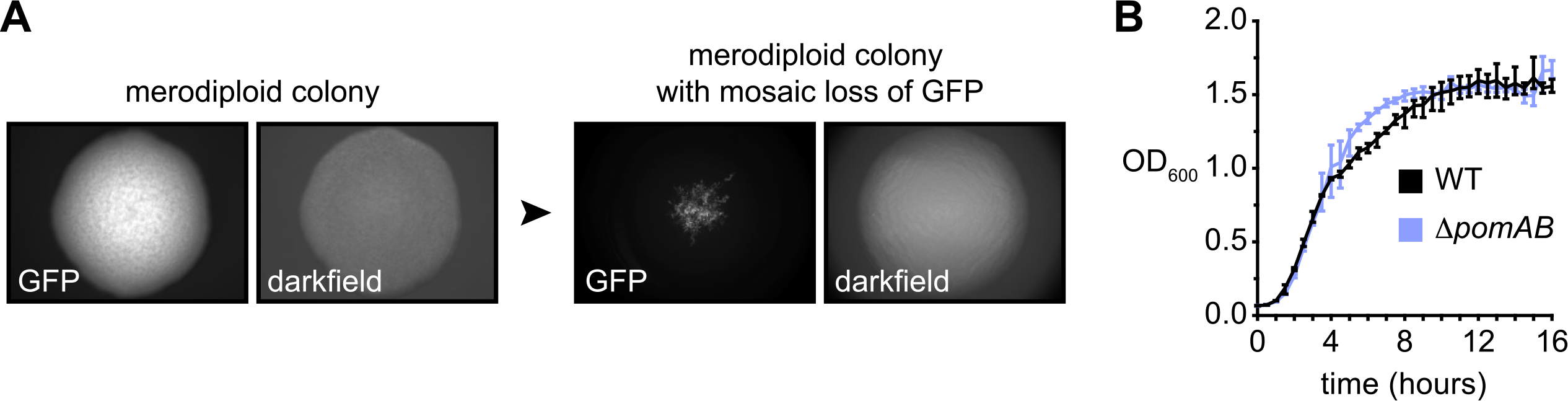
Additional information related to the engineering of the *Vibrio* ZWU0020 Δ*pomAB* deletion mutant. (**A**) *Vibrio* ZWU0020 *pomAB* merodiploids robustly express GFP. After outgrowth and plating on nonselective media, mosaic merodiploid colonies exhibiting loss of GFP and thus, loss of the allelic exchange vector, can be isolated. Images labeled “GFP” show GFP fluorescence and “darkfield” show total colony structure. (**B**) Plotted is the average optical density at 600nm (OD_600_) vs. time (hours) of WT ZWU0020 and its Δ*pomAB* derivative during shaking growth in LB broth at 30°C. Range bars are based on four technical replicates.

**Figure 6-Figure Supplement 2.**
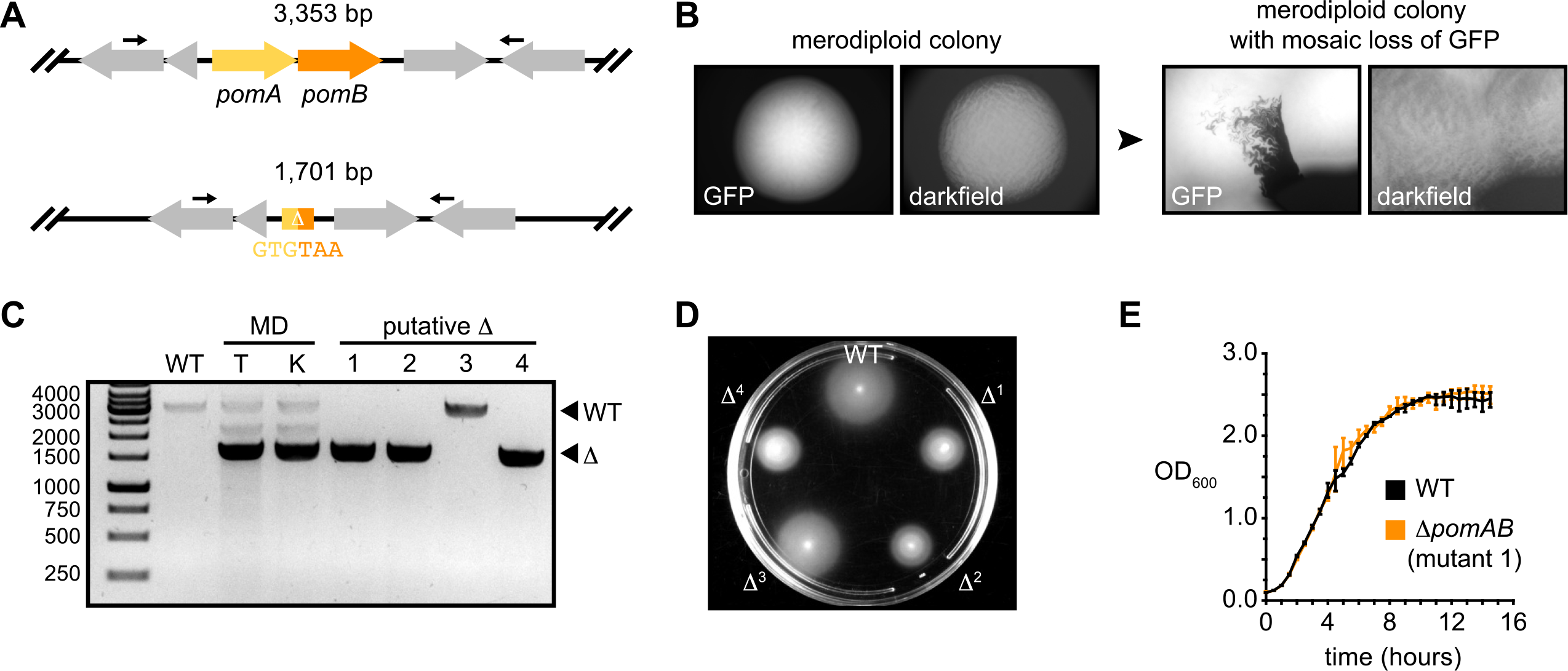
Markerless deletion of *pomAB* in *Aeromonas veronii* using the allelic exchange vector pAX2. (**A**) Top: wild-type *pomAB* locus in *Aeromonas veronii* ZOR0001. Bottom: result of markerless *pomAB* deletion fuses the start codon of *pomA* to the stop codon of *pomB*. Black arrows mark approximate primer annealing sites for genotyping and the size of each amplification product is indicated. (**B**) *Aeromonas* ZOR0001 *pomAB* merodiploids robustly express GFP. After outgrowth and plating on nonselective media, mosaic merodiploid colonies exhibiting loss of GFP, and thus loss of the allelic exchange vector, can be isolated. Images labeled “GFP” show GFP fluorescence and “darkfield” show total colony structure. (**C**) Agarose gel showing PCR-based genotyping of wild-type (WT), merodiploid (MD), and four putative Δ*pomAB*(Δ) mutants. To verify the dual functionality of pAX2, two merodiploids were isolated using either its temperature (T) or kill switch (K) donor cell counterselection systems. Putative Δ*pomAB* mutants were derived from “K” merodiploids. Migration distances of WT and mutant alleles are indicated. Mutant 3 is an example of a wild-type revertant, whereas 1, 2, and 4 are Δ*pomAB* mutants. (**D**) Swim motility of WT ZOR0001 and the four putative Δ*pomAB* mutants from **C** in 0.2% tryptic soy agar at 30°C. The attenuated swimming phenotype of mutants 1, 2, and 4 corroborates genotyping results. Of note, *Aeromonas* ZOR0001 encodes multiple lateral flagellar systems in addition to one polar flagellum; therefore, partial loss of motility is expected. (**E**) Plotted is the average optical density at 600nm (OD_600_) vs. time (hours) of WT ZOR0001 and Δ*pomAB* mutant 1 during shaking growth in LB broth at 30°C. Range bars are based on four technical replicates.

**Figure 6-Figure Supplement 3.**
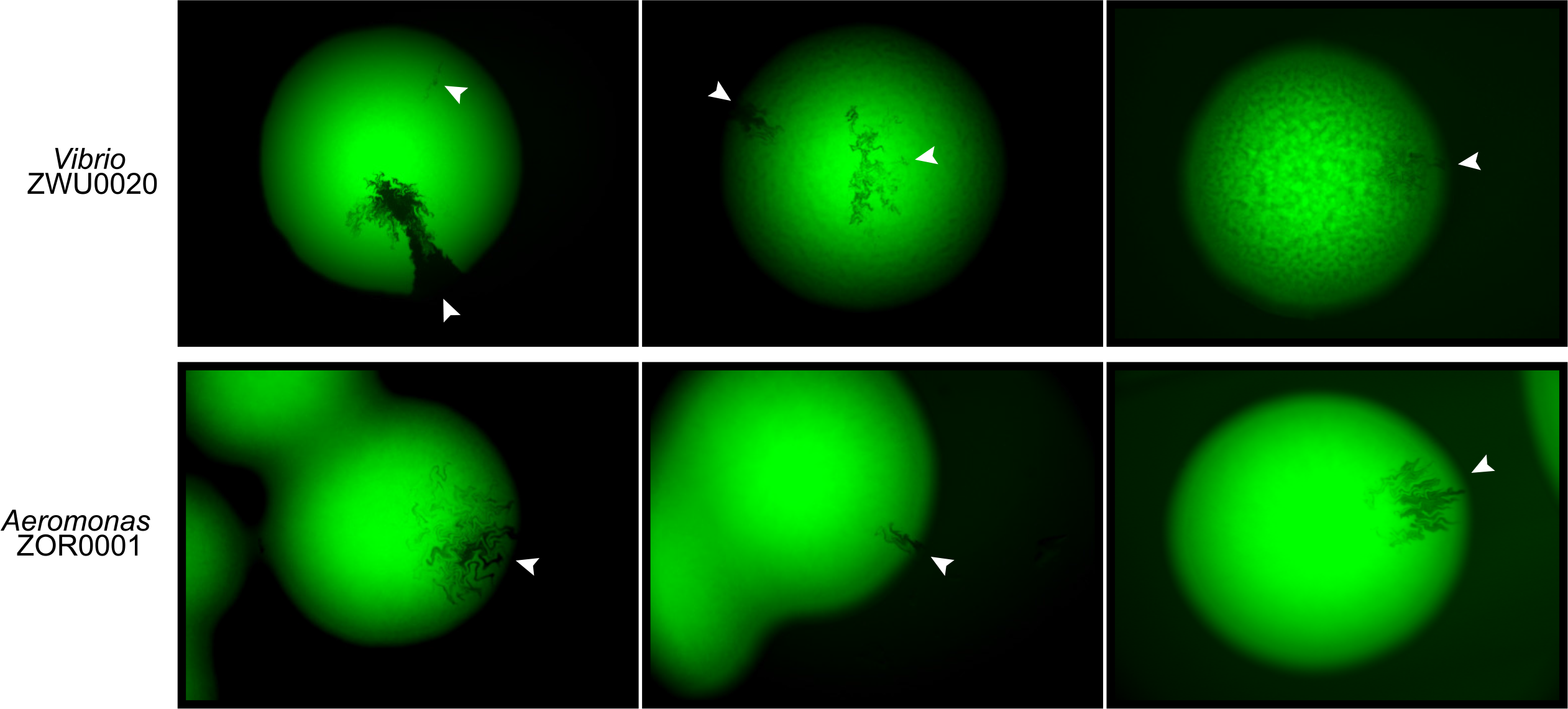
Examples of various patterns of GFP loss observed within merodiploid colonies. *Vibrio* ZWU0020 (top row) and *Aeromonas* ZOR0001 (bottom row) merodiploid cells were plated on non-selective tryptic soy agar. After overnight growth, colonies were screened for loss of GFP, which indicates that a second recombination event has occurred. White arrowheads mark GFP-negative patches.

**Figure 7-Figure Supplement 1.**
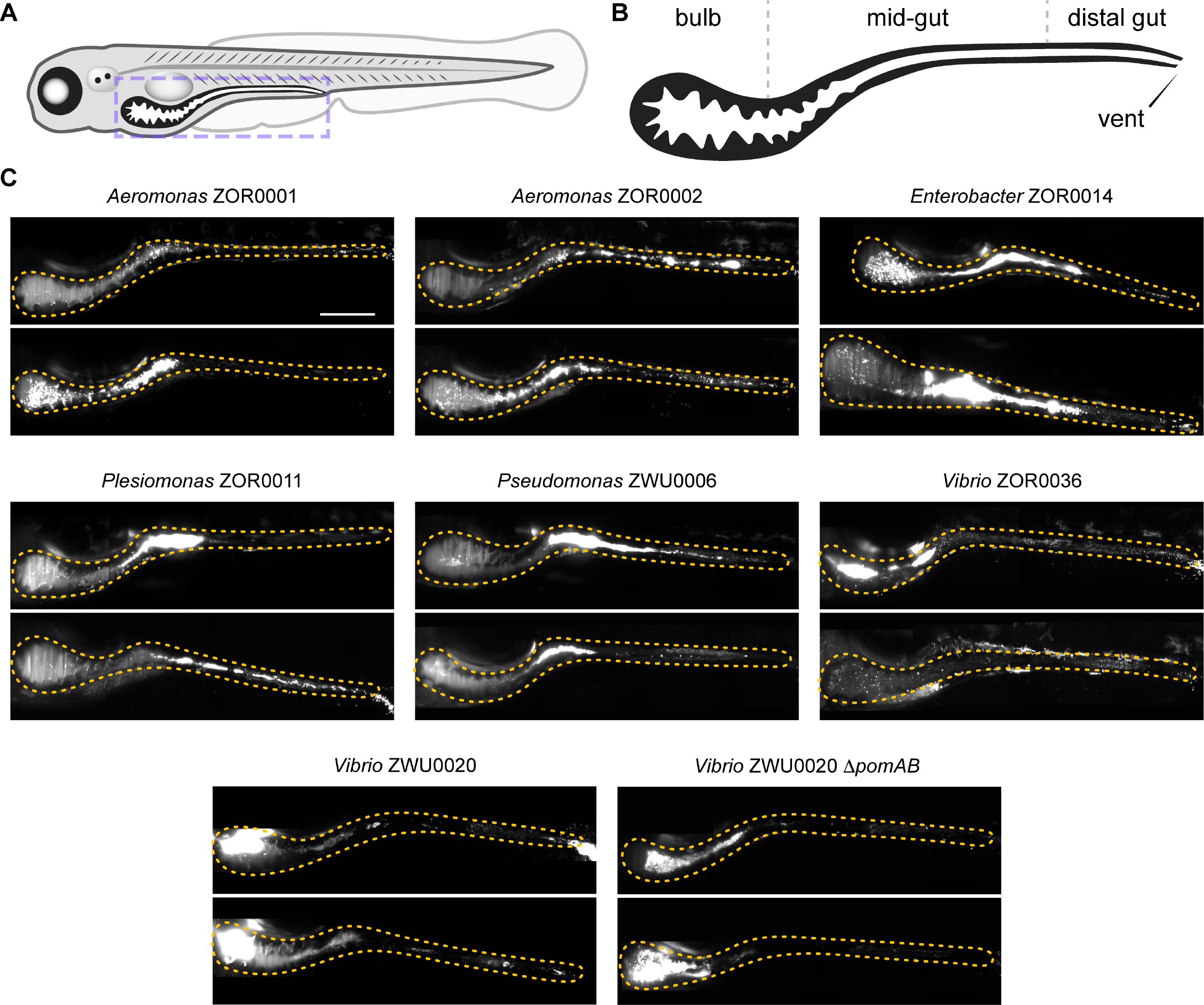
Additional images showing intestinal colonization patterns and growth modes of zebrafish symbionts. (**A**) Cartoon diagram of a 5-day old larval zebrafish. Purple dashed box outlines region imaged in **C**. (**B**) Diagram shows the boundaries of the bulb, mid-gut, and distal gut within the larval intestine. (**C**) Maximum intensity projections of 3D image stacks acquired by light sheet fluorescence microscopy for indicated bacterial strains. Orange dotted outline marks the intestine in each image. Scale bar: 200μm.

**Figure 7-Figure Supplement 2.**
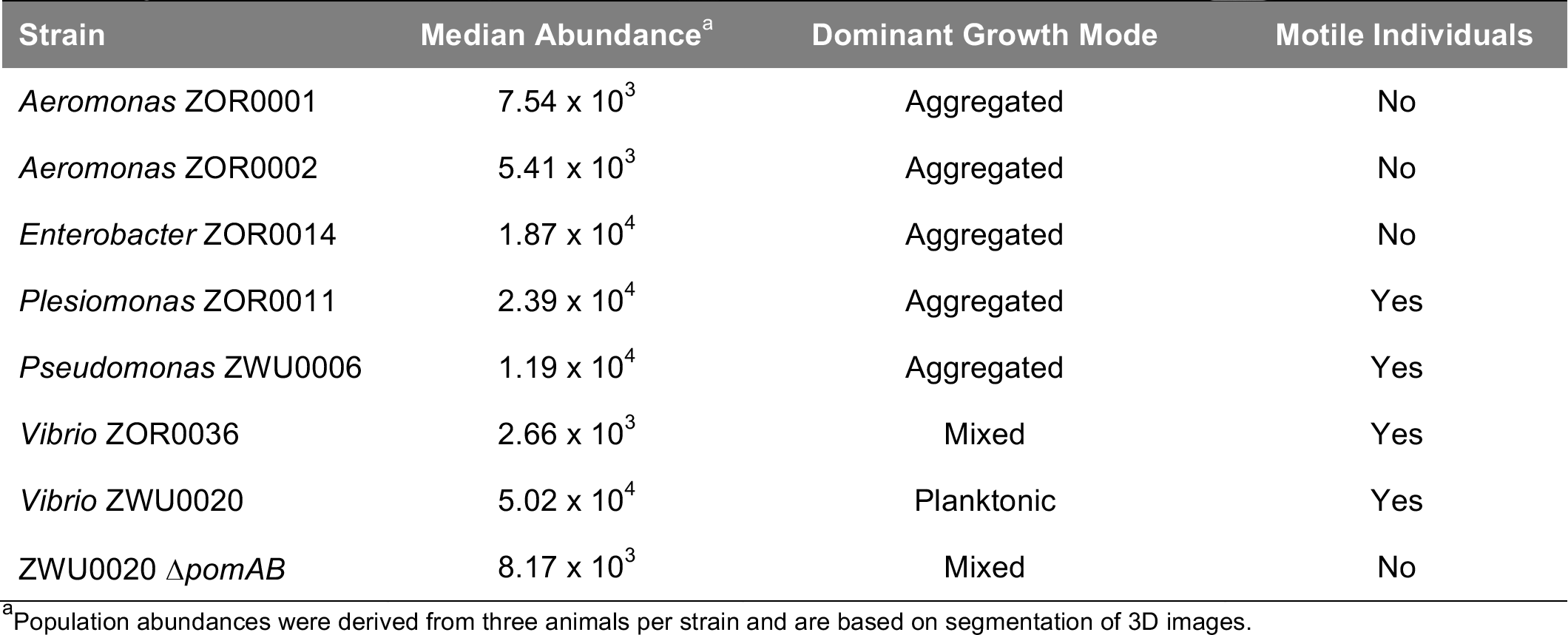
Summary of traits exhibited by bacterial symbionts within the larval zebrafish intestine.

**Figure 7-Figure Supplement 3.**
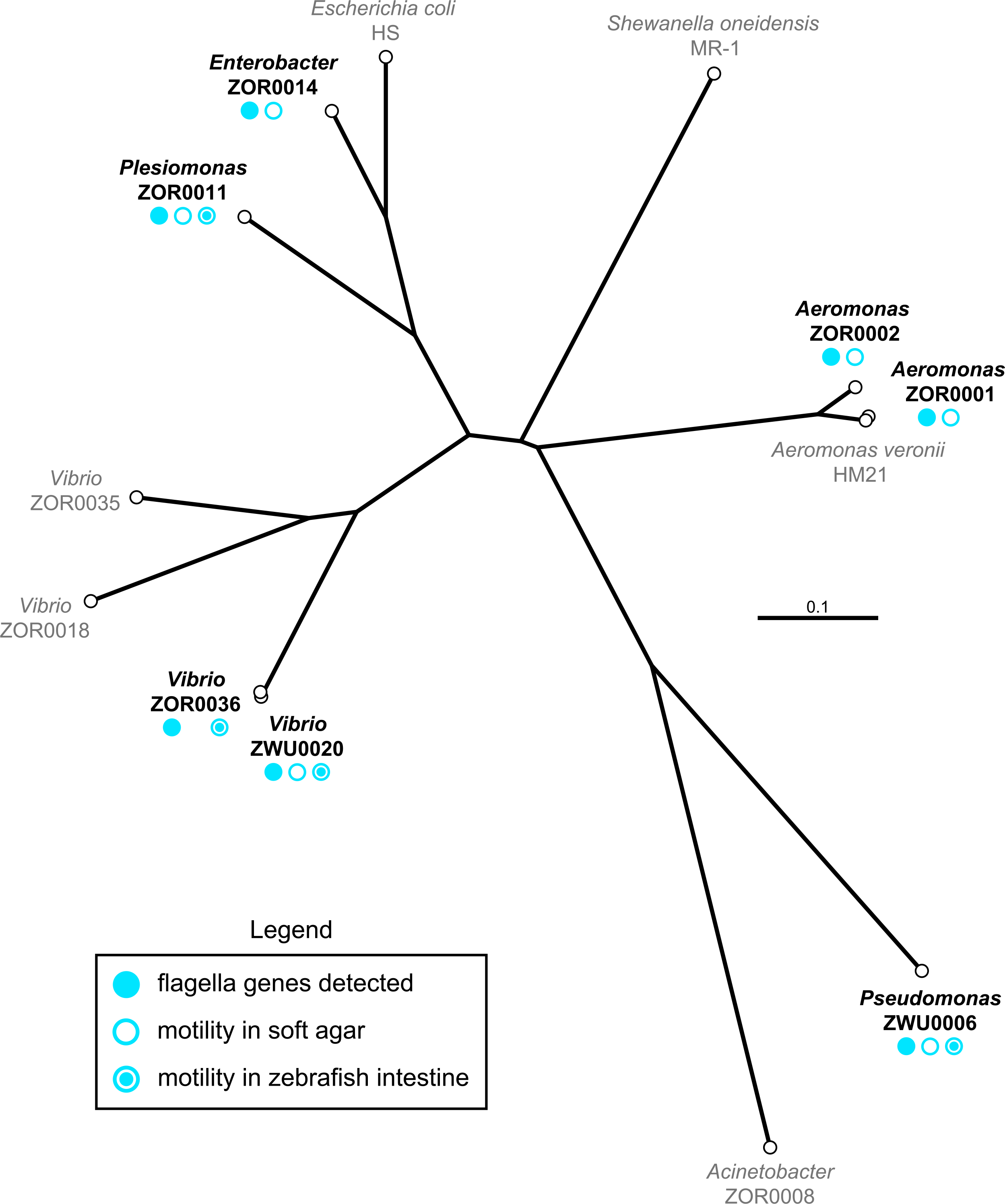
Phylogenetic relatedness and summary of motility phenotypes. Shown is an unrooted phylogenetic tree generated using full-length nucleotide sequences of the 16S rRNA gene from all strains manipulated in this study. Strains used for live imaging are in bold black type and symbols denote motility phenotypes.

**Figure 7-Figure Supplement 4.**
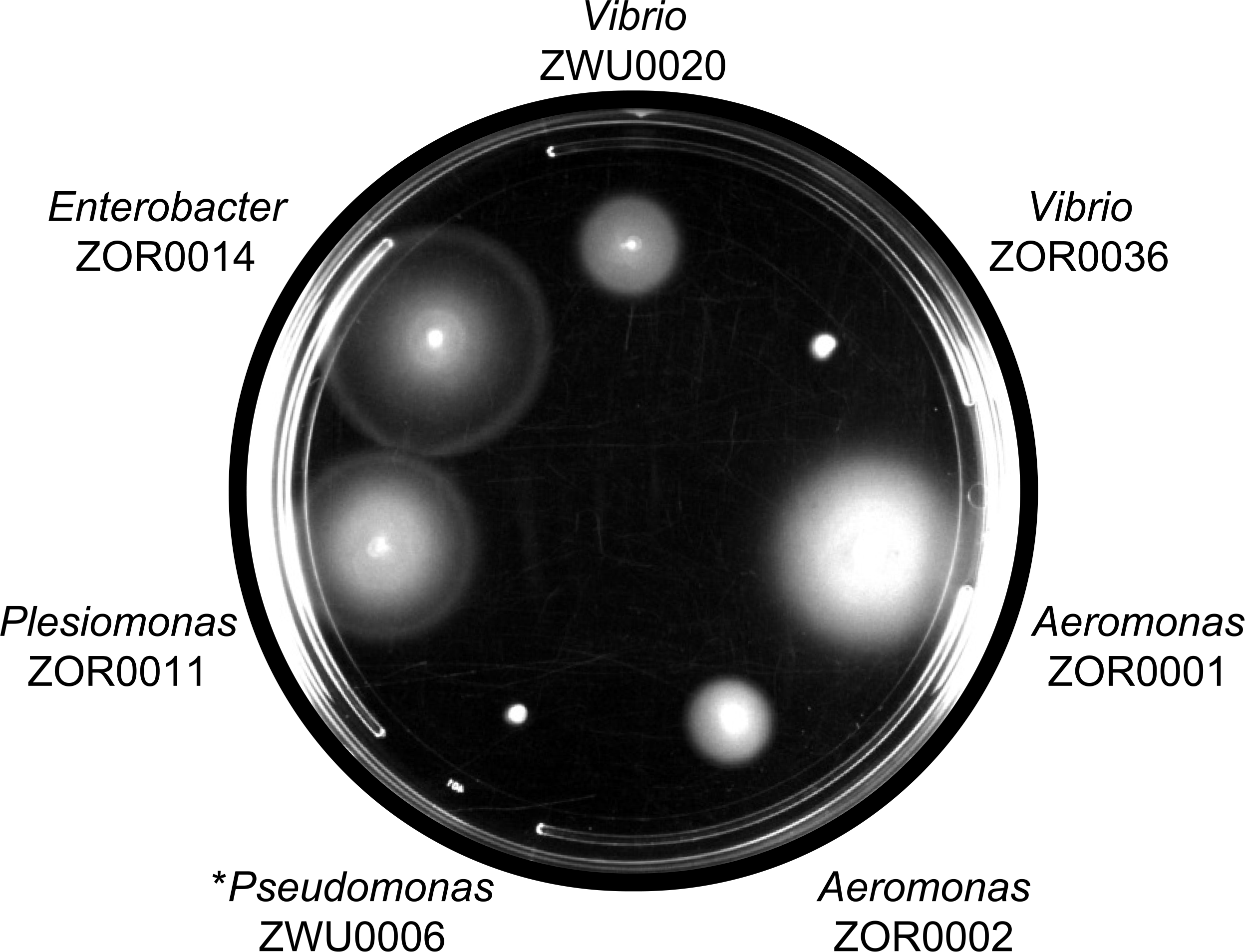
Motility of select strains in soft agar. Indicated strains were inoculated in 0.2% tryptic soy agar and incubated at 30°C for ~5h. * *Pseudomonas* ZWU0006 exhibits a delay in swimming and advances after a 24h incubation period.

**Supplementary Movie 1.** Example of *Vibrio* ZWU0020 growth mode and behavior within the zebrafish gut. Movie depicts live imaging of a single optical plane in the intestinal bulb of a 5-day old larval zebrafish colonized with a 100:1 mixture of *Vibrio* ZWU0020 expressing dTomato (left panel) or sfGFP (right panel). The GFP-tagged subpopulation highlights the highly motile and planktonic nature of this strain within the intestine. Scale bars: 50μm.

**Supplementary Movie 2.** Example of *Enterobacter* ZOR0014 growth mode and behavior within the zebrafish gut. Movie depicts live imaging of a single optical plane in the intestinal bulb of a 5-day old larval zebrafish colonized with *Enterobacter* ZOR0014 expressing sfGFP. A small portion of the *Enterobacter* mid-gut population can be seen fluxing into the bulb due to peristaltic contractions. Non-motile cells are evident along with larger multicellular aggregates. Scale bar: 50μm.

**Supplementary Movie 3.** Example of *Aeromonas* ZOR0001 growth mode and behavior within the zebrafish gut. Movie depicts live imaging of a single optical plane in the intestinal mid-gut of a 5-day old larval zebrafish colonized with *Aeromonas* ZOR0001 expressing sfGFP. Non-motile cells and small aggregates can be seen experiencing flux due to peristaltic contractions. Scale bar: 50μm.

**Supplementary Movie 4.** Example of *Aeromonas* ZOR0002 growth mode and behavior within the zebrafish gut. Movie depicts live imaging of a single optical plane in the intestinal bulb of a 5-day old larval zebrafish colonized with *Aeromonas* ZOR0002 expressing sfGFP. Non-motile cells and small aggregates can be observed. Scale bar: 50μm.

**Supplementary Movie 5.** Example of *Vibrio* ZOR0036 growth mode and behavior within the zebrafish gut. Movie depicts live imaging of a single optical plane in the intestinal bulb of a 5- day old larval zebrafish colonized with *Vibrio* ZOR0036 expressing sfGFP. Highly motile cells as well as large multicellular aggregates can be observed. Scale bar: 50μm.

**Supplementary Movie 6.** Example of *Plesiomonas* ZOR0011 growth mode and behavior within the zebrafish gut. Movie depicts live imaging of a single optical plane in the intestinal bulb of a 5-day old larval zebrafish colonized with *Plesiomonas* ZOR0011 expressing sfGFP. Motile cells as well as large multicellular aggregates can be observed. Notably, the swim speed of *Plesiomonas* cells appears to be more moderate compared to *Vibrio* or *Pseudomonas* strains. Scale bar: 50μm.

**Supplementary Movie 7.** Example of *Pseudomonas* ZWU0006 growth mode and behavior within the zebrafish gut. Movie depicts live imaging of a single optical plane in the intestinal bulb of a 5-day old larval zebrafish colonized with *Pseudomonas* ZWU0006 expressing sfGFP. Highly motile cells as well as large multicellular aggregates can be observed. Scale bar: 50μm.

**Supplementary Movie 8.** Example of *Vibrio* ZWU0020 Δ*pomAB* growth mode and behavior within the zebrafish gut. Movie depicts live imaging of a single optical plane in the intestinal bulb of a 5-day old larval zebrafish colonized with *Vibrio* ZWU0020 Δ*pomAB* expressing dTomato. Non-motile cells and small aggregates can be observed. Scale bar: 50μm.

### SUPPLEMENTARY AND SOURCE DATA FILE LEGENDS

**Supplementary File 1.** Wild and Recombinant Bacteria.

**Supplementary File 2.** 16S rRNA Nucleotide Sequences.

**Supplementary File 3.** *E. coli* Strains and Plasmids.

**Supplementary File 4.** Primers.

**Supplementary File 5.** Nucleotide Sequences of Select Genetic Parts.

**Supplementary File 6.** Plasmid Construction.

**Supplementary File 7.** Protocol: Tn7 tagging with pTn7xTS and pTn7xKS

**Supplementary File 8.** Protocol: Allelic exchange with pAX1 and pAX2

**Figure 1-Source Data 1.** Raw in vitro growth data for Figure 1A

**Figure 3-Source Data 1.** Raw in vitro growth data for Figure 3-Figure Supplement 2C

**Figure 6-Source Data 1.** Raw in vitro growth data for Figure 6-Figure Supplement 1B

**Figure 6-Source Data 2.** Raw in vitro growth data for Figure 6-Figure Supplement 2E

**Figure 8-Source Data 1.** Values plotted in Figure 8

